# Hello darkness, my old friend: 3-Ketoacyl-Coenzyme A Synthase4 is a branch point in the regulation of triacylglycerol synthesis in *Arabidopsis thaliana*

**DOI:** 10.1101/2020.07.27.223388

**Authors:** Urszula Luzarowska, Anne-Kathrin Ruß, Jérôme Joubès, Marguerite Batsale, Jędrzej Szymański, Venkatesh Periyakavanam Thirumalaikumar, Marcin Luzarowski, Si Wu, Feng Zhu, Niklas Endres, Sarah Khedhayir, Julia Schumacher, Weronika Jasinska, Ke Xu, Sandra Marcela Correa Cordoba, Simy Weil, Aleksandra Skirycz, Alisdair Robert Fernie, Yonghua Li-Beisson, Corina Mariana Fusari, Yariv Brotman

**Affiliations:** Department of Life Sciences, Ben Gurion University of the Negev, Beersheva, Israel; Max Planck Institute of Molecular Plant Physiology, Am Mühlenberg 1, 14476 Potsdam-Golm, Germany; University Bordeaux, CNRS, Laboratoire de Biogenèse Membranaire, UMR 5200, F-33140 Villenave d’Ornon, France; Department of Molecular Genetics, Leibniz Institute of Plant Genetics and Crop Plant Research, OT Gatersleben, 06466, Seeland, Germany; Department of Genetics, Stanford University School of Medicine, Stanford, CA 94305, USA; National R&D Center for Citrus Preservation; Key Laboratory of Horticultural Plant Biology, Ministry of Education; Huazhong Agricultural University, Wuhan 430070, P.R. China; Department for Plant Cell and Molecular Biology, Institute for Biology, Humboldt-Universität zu Berlin, Philippstraße 13, 10115 Berlin, Germany; Aix Marseille Univ., CEA, CNRS, BIAM, Institute de Biosciences et Biotechnologies Aix-Marseille, F-13108 Saint Paul-Lez-Durance, France; Centro de Estudios Fotosintéticos y Bioquímicos (CEFOBI-CONICET-UNR), Suipacha 570, S2000LRJ Rosario, Argentina

**Keywords:** Arabidopsis, GWAS, natural variation, triacylglycerols, fatty acids, KCS4, carbon starvation

## Abstract

Due to their sessile lifestyle, plants have evolved unique mechanisms to deal with environmental challenges. Under stress, plant lipids are important as alternative sources of carbon and energy when sugars or starch are limited. Here, we applied combined heat and darkness and extended darkness to a panel of ∼ 300 Arabidopsis accessions to study lipid remodeling under carbon starvation. Natural allelic variation at *3-KETOACYL-COENZYME A SYNTHASE4* (*KCS4*), a gene encoding for an enzyme involved in very long chain fatty-acid (VLCFA) synthesis, underlies a differential accumulation of polyunsaturated triacylglycerols (puTAGs) under stress. Ectopic expression in yeast and plants proved that KCS4 is a functional enzyme localized in the ER with specificity for C22 and C24 saturated acyl-CoA. Allelic mutants and transient overexpression *in planta* revealed the differential role of *KCS4* alleles in VLCFA synthesis and wax coverage, puTAG accumulation and biomass. Moreover, the region harboring *KCS4* is under high selective pressure and allelic variation at *KCS4* correlated with environmental parameters from the locales of Arabidopsis accessions. Our results provide evidence that KCS4 plays a decisive role in the subsequent fate of fatty acids released from chloroplast-membrane lipids under carbon starvation. This work sheds light on both plant response mechanisms and the evolutionary events shaping the lipidome under carbon starvation.

**One sentence summary:** Natural variation at *KCS4* underlies a differential accumulation of polyunsaturated triacylglycerols under carbon starvation, by acting as a regulatory branch point in the fate of fatty acids.

## Introduction

Plant metabolism profoundly influences or even co-ordinates signaling, physiological, and defense responses. When the prevailing environment is adverse, biosynthesis, concentration, transport, and storage of primary and secondary metabolites are affected, in a stress dependent manner. These metabolic responses to abiotic stress involve fine adjustments in carbohydrate, amino acid, amine and lipid pathways. Proper activation of early metabolic responses helps cells restore chemical and energetic imbalances imposed by the stress and is crucial to acclimation and survival (Fraire-Velázquez and Balderas-Hernández, 2013).

Like many other complex traits, plant metabolism has an extended variation within natural populations, and therefore has been widely investigated by using genome-wide association studies (GWAS). Genetic factors involved in variation for key metabolic traits have been successfully identified and characterized (i.e., primary and secondary metabolites and enzyme activities) (Wu et al., 2016, 2018; Fusari et al., 2017). A number of GWAS regarding lipid metabolism have been reported and resulted in the identification of new enzymes and regulators in species like Arabidopsis, maize, coffee, rice and canola (Riedelsheimer et al., 2013; Gacek et al., 2017; Sant’Ana et al., 2018; Li et al., 2019; Hong et al., 2022). These studies focused on lipid levels under optimal growth conditions. However, accumulation of lipids is highly affected by fluctuations in the environmental conditions, such as light, temperature and nutrient availability (Burgos et al., 2011; Caldana et al., 2011; Szymanski et al., 2014). A classic example involves compensation for the decreased membrane fluidity under cold stress (Welti et al., 2002). Moreover, adaptation to a high temperature has been associated with an increase in the relative content of digalactosyldiacylglycerol (DGDG) and the ratio of DGDG to monogalactosyldiacylglycerol (MGDG), as well as a moderate increase in fatty-acyl saturation of plant lipids (Han, 2016).

In plants, the major source of energy in the dark is starch, which is degraded linearly according to the duration of the previous night (Graf and Smith, 2011). Therefore, plants subjected to a sudden extended night (dark stress) experience carbon starvation due to limited starch availability. Under these conditions, polyunsaturated fatty acids (PUFA) released from membrane lipids are used as an alternative substrate for respiration via β-oxidation. These released fatty acids are often sequestered into triacylglycerols (TAGs) to avoid the adverse effects of the detergent-like properties of free fatty acids (FFAs). Therefore, it follows that TAG synthesis and lipid-droplet formation sometimes constitute a cellular response to lipotoxicity. This sequestration is especially true for PUFA species because they contain multiple double bonds and are liable to production of reactive oxygen species (ROS) (Kao et al., 2018). It has been shown that the process of FFA sequestration into TAGs protects the cell from ROS generated during PUFA peroxidation (Fan et al., 2017).

Major changes in lipidomic composition under stress are accompanied by a remarkable remodeling of gene expression, both at the level of whole metabolic pathways, but also involving activation of specific stress-induced isozymes (Szymanski et al., 2014). This work suggests that only by causing a perturbation of the metabolic homeostasis is GWAS likely to capture genes that evolved to stand at critical branch points in the regulation of the lipidome under natural fluctuating environments.

Here, we performed GWAS using lipidomic data obtained in four different environments: control, combined heat and darkness, and two different extended darkness conditions. The locus harboring *KCS4*, encoding for a *3-KETOACYL-COENZYME A SYNTHASE* (Joubès et al., 2008; Kim et al., 2021), was strongly associated with a high number of polyunsaturated TAGs (puTAGs) in stress conditions, but was not detected in control condition. Thus, *KCS4* association is driven by carbon starvation. Using mutants for alternative *KCS4* alleles and ectopic expression in yeast and plants we explored the biochemical properties of *KCS4* and validated its role in lipid metabolism. We place a model for *KCS4* enzymatic and regulatory function, where *KCS4* acts as a branch point in the fate of FAs, directing saturated FA to the very-long chain fatty acid pathway (VLCFA) and facilitating the pool of polyunsaturated FA in the accumulation of puTAGs. Two *KCS4* alleles have differential capacity to process saturated FA. Accessions carrying the strong allele show higher levels of VLCFA-derived cuticular waxes and higher levels of puTAGs. In fully exploiting the genetic data available for Arabidopsis, this work sheds light on the evolutionary events shaping the lipidome and further highlights the importance of lipid remodeling in plant response to changing environments.

## Results

### Plasticity in Arabidopsis lipid metabolism under different environmental conditions

In order to investigate the genetic regulation of lipid biosynthesis in Arabidopsis, we conducted LC– MS-based lipidomic analysis of 301 and 309 Arabidopsis accessions grown in two environmental conditions: control (21/16°C in 16/8h light/dark cycle, termed CHD) and stress (heat 32°C + darkness for 24 h, termed HD), respectively. HD was selected as it has previously been shown to have a strong effect on the lipidome (Caldana et al., 2011). The mapping population was phenotyped for a total of 98 and 109 glycerolipids in CHD and HD, respectively (Supplementary Dataset 1).

Principal component analysis (PCA) of lipid data clearly separates the natural ecotypes into two different groups according to the two environmental conditions tested (Supplementary Figure 1a). Furthermore, differences in the lipid status between CHD and HD samples showed that TAGs were strongly accumulated while MGDGs and DGDGs were mostly decreased under stress (Supplementary Figure 1b). In addition, there is substantial positive correlation between lipid levels from a given lipid class, suggesting coordinated regulation at the level of lipid classes (Supplementary Figure 1c). Notably, TAGs showed the highest variation across accessions for both stress and control conditions, with many species falling in the range of >10-fold difference (Supplementary Figure 1d).

**Figure 1.**
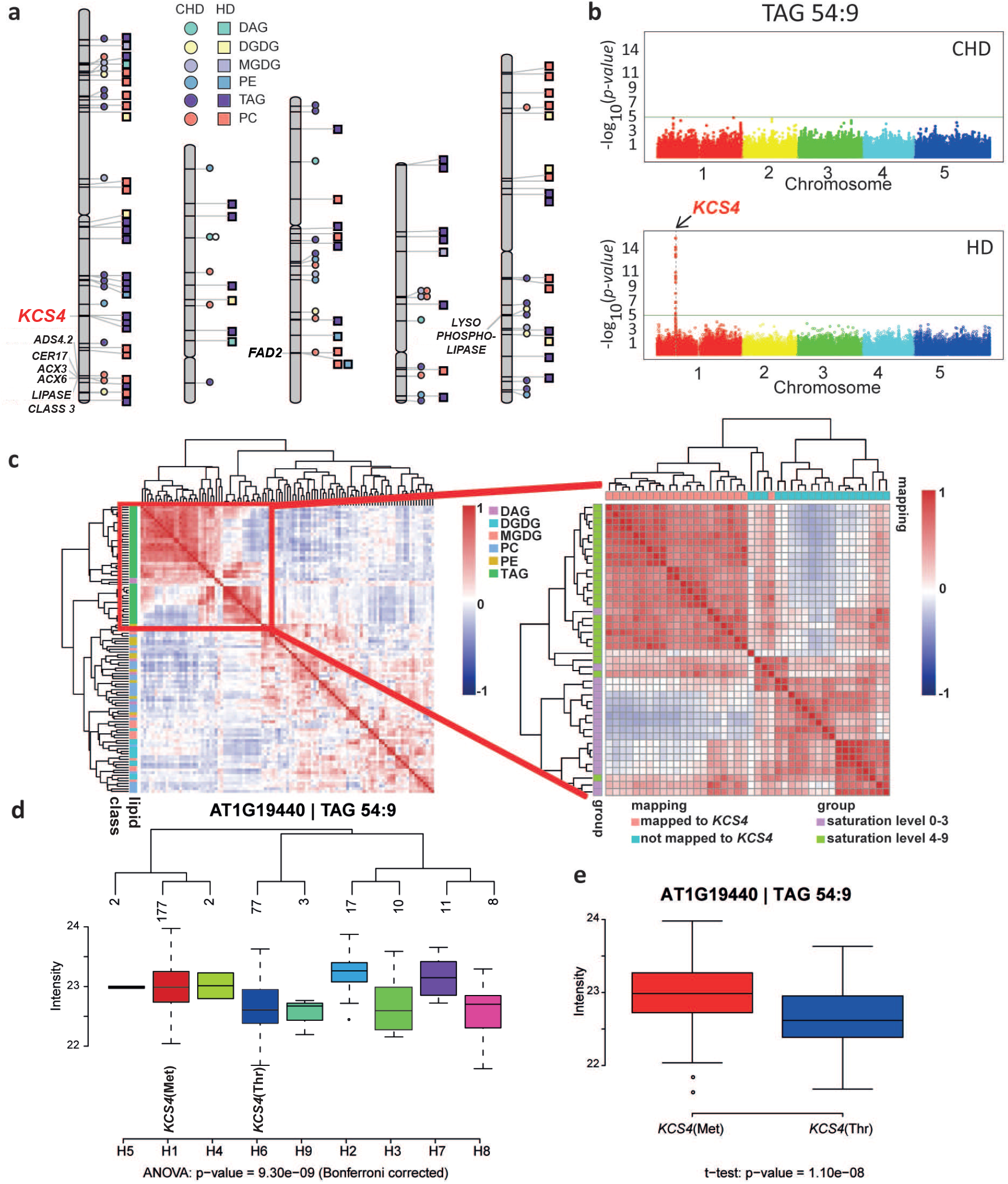
Results from GWAS analysis on lipidomic data from plants grown under control (CHD) and stress (HD) conditions. **a**. Chromosome scheme and Quantitative Trait Loci (QTL) identified in the two experimental conditions (CHD = circles, HD = squares), for different lipid classes: phosphatidylethanolamine (PE), phosphatidylcholine (lecithin) (PC), monogalactosyldiacylglycerol (MGDG), digalactosyldiacylglycerol (DGDG), diacylglycerol (DAG) and triacylglycerol (TAG). Color/shape references are included in the figure. Gene names are included only for co-localizing QTL for two or more lipid species with LOD > 5.3. **b**. Manhattan plots obtained for TAG 54:9 in CHD (top) and in HD (bottom). The *KCS4* QTL was found at Chromosome 1 only under HD condition. **c**. Pearson correlations coefficients calculated between all lipid species across all accessions, for HD condition. Lipids are color-coded according to class (see reference at right) and clustered using Ward’s distance on pairwise dissimilarity. A zoom-in of the TAG cluster shows the grouping pattern of TAGs according to number of unsaturated bonds and to the GWAS result (left). **d**. Average trait value (intensity of TAG 54:9, log2 scale) for the different *KCS4* haplotypes using SNPs m11502, m11503, m11504 and m11505. The two major haplotypes have the alternative alleles for m11503 (Met or Thr) **e**. TAG 54:9 average value for accessions carrying the *KCS4*(Met) and *KCS4*(Thr) alleles.

### Genetic basis underlying Arabidopsis lipid metabolism

Lipid intensities from CHD and HD and the genotypic information from the 250K SNPchip (Horton et al., 2012) were used to perform two independent genome-wide association studies (GWAS) for the identification of lipid quantitative trait loci (QTL). GWAS revealed 93 and 544 SNP–trait associations in control and stress conditions, respectively, at a genome-wide significance level of LOD ≥ 5.3 (Supplementary Dataset 2). These SNP–trait associations further grouped into 51 and 66 QTL for CHD and HD conditions, respectively. In total, only five QTL were shared between CHD and HD, revealing the great plasticity of the lipidome in response to different environmental conditions (Supplementary Dataset 3).

Regarding functional annotation of candidate genes, we mapped 12 and 22 genes putatively involved in lipid metabolism for control and stress condition, respectively (Supplementary Dataset 3, Acyl-Lipid Metabolism database (Li-Beisson et al., 2010) http://aralip.plantbiology.msu.edu/) (Figure 1). After filtering for robust associations co-localizing for two or more traits, four QTL harbored lipid-genes (Figure 1a). Two QTL in Chromosome 1, one harboring the *3-KETOACYL-COENZYME A SYNTHASE4* (*KCS4*, AT1G19440) and other including a *LIPASE CLASS 3* (AT1G06250), two *ACYL-COA OXIDASES* (*ACX3*: AT1G06290 and *ACX6*: AT1G06310) and two *FATTY ACID DESATURASE* (*CER17*: AT1G06350, ADS4.2: AT1G063609); one in Chromosome 3 harboring the *FATTY ACID DESATURASE* (*FAD2:* AT3G12120) and the last QTL in Chromosome 5, harboring a *LYSOPHOSPHOLIPASE* (AT5G20060) (Supplementary Dataset 3, Figure 1a).

### *3-KETOACYL-COENZYME A SYNTHASE4* is involved in fatty acid elongation and has major impact on polyunsaturated TAG levels under heat and dark stress

Remarkably, 22 polyunsaturated TAGs (puTAGs) co-localized to the QTL on chromosome 1, harboring *KCS4* (Figure 1a). This QTL was not found in control conditions, and the lead SNP in the locus (m11481) co-localized with LOD ≥ 8 (Bonferroni corrected p-value < 0.05) for 16 of the 22 associated TAGs (e.g., TAG 54:9, Figure 1b; Supplementary Dataset 2). Besides *KCS4*, 33 other genes mapped on the QTL, spanning 107,318 bp (Supplementary Dataset 3). However, with the exception of *KCS4*, none of the others showed any homology to proteins known to be involved in lipid metabolism.

*KCS4* belongs to a gene family involved in the elongation of VLCFA. Twenty-one *KCS* genes have been previously identified in the Arabidopsis genome. The *KCS* family encodes for enzymes involved in the first and often described as rate-limiting step of fatty–acid elongation (FAE) (Blacklock and Jaworski, 2006; Joubès et al., 2008). KCS4’s closest homologues are: KCS9 (78% aa identity), involved in the elongation of C22 to C24 acyl-CoAs (Kim et al., 2013); KCS17 (72% aa identity); and KCS18 (64% aa identity), involved in elongating C18 acyl-CoAs to C20 and C22 in seeds and shown to drive the natural variation in the abundance of VLCFAs in this organ (Rossak et al., 2001). However, KCS4 specificity and function have not been clearly characterized so far. TAGs associated with the *KCS4* QTL were exclusively polyunsaturated (e.g., puTAG 52:6, 52:7, 54:6, 54:8, 54:9, 56:6, Supplementary. Table 2-3) and correlated positively with each other, grouping together in a single well-defined cluster (Figure 1c).

To identify the alleles involved in puTAG variation under HD, we constructed haplotypes considering the four SNPs from *KCS4* scored in the 250K SNPchip (m11502, m11503, m11504, m11505, Supplementary Dataset 2). The population divided into nine non-unique haplotypes, which were therefore, further considered as being putatively involved in the QTL variation. Mean levels of puTAG 54:9 varied significantly among haplotypes (Figure 1d). Furthermore, m11503 was identified as a non-synonymous SNP (nsSNP). In the majority of accessions, SNP m11503 encodes for a methionine [hereafter *KCS4*(Met)]. Accessions carrying this allele have higher levels of puTAG in the stress condition than accessions carrying the minor allele, which encodes for a threonine [hereafter *KCS4*(Thr)] (Figure 1e). These results suggest that the nsSNP (m11503) in *KCS4* is the causal mutation underlying puTAG variation under HD conditions in Arabidopsis.

### *KCS4* associates with polyunsaturated TAGs in extended darkness

Thus far, we found that *KCS4* associates with puTAGs in a combined heat and darkness treatment. In prolonged darkness, starch is prematurely exhausted and plants are carbon starved (Stitt and Zeeman, 2012). Therefore, we determined whether *KCS4* also associates with puTAGs, when darkness alone is applied.

For this purpose, we utilized own data sets: control (CD), 3 days (3D) extended darkness, and 6 days (6D) extended darkness (Supplementary Dataset 1) (Zhu et al., 2022). The GWAS at a genome-wide significance level of LOD ≥ 5.3 resulted in 230, 255 and 342 SNP–trait associations grouped in 73, 81, and 70 QTL for CD, 3D, and 6D, respectively (Supplementary Dataset 2, Supplementary Dataset 3). The locus harboring *KCS4* associated strongly in 3D and 6D with 8 and 13 puTAGs, respectively. Lead SNP and other features of the association matched with GWAS results from the HD experiment (Supplementary Figure 2). Thus, darkness alone is enough to trigger the association of *KCS4* with puTAGs and could be related to the carbon status of the plant. Expression of *DIN1* (AT4G35770) and *BCAT2* (AT1G10070), two genes usually used as starvation markers (Moraes et al., 2019) are upregulated under HD conditions (Supplementary Figure 3), showing that HD is also triggering carbon starvation. In summary, an additive effect is observed when both stresses are applied together, determining a stronger association of *KCS4* to puTAGs under HD.

**Figure 2.**
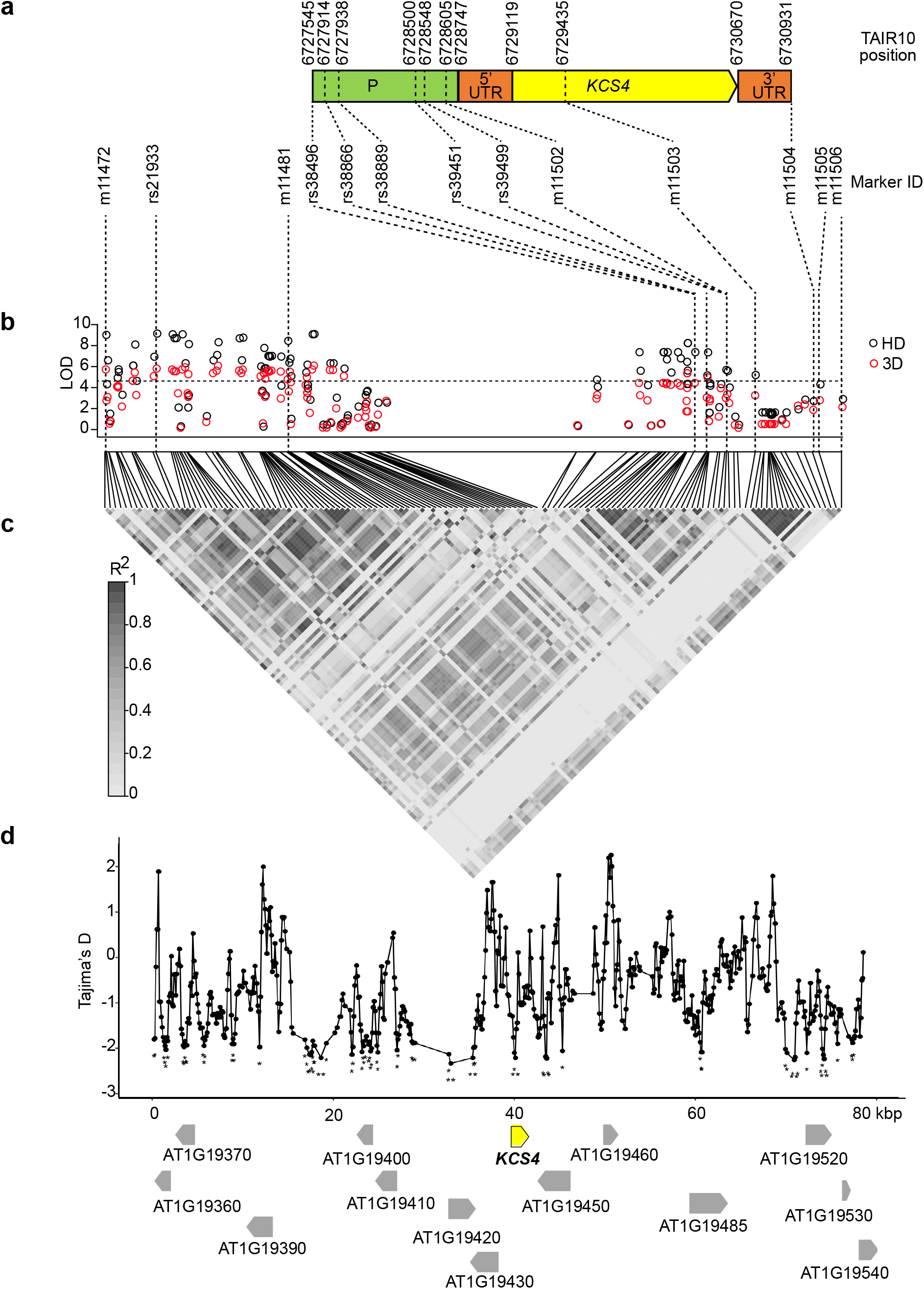
Dissection of *KCS4* alleles involved in TAG variation, pattern of linkage disequilibrium and signatures of selection. **a**. *KCS4* gene model depicting promoter region (P, green), 5’UTR (orange), coding region (yellow) and 3’UTR (orange) of AT1G19440/*KCS4*. Beginning, end and relevant SNP positions are indicated above (TAIR10) and bellow (SNP ID) gene structure. **b**. Manhattan plot of GWAS results (HD: black circles; 3D: red circles) obtained for TAG 54:9 using re-sequenced information for 149 accessions. The re-sequenced region on Chromosome 1 (positions 6689119 to 6769118, TAIR10) included 131 SNPs genotyped in the 250K SNPchip plus 1,137 additional polymorphisms. Bonferroni threshold (LOD = 4.4) is indicated with a horizontal dashed line. **c**. Linkage disequilibrium (LD) is depicted as a heat map of the coefficient of correlation (r^2^). The LD block (0.8 > r^2^ > 0.2, P < 0.001) includes the lead SNPs rs21933 and m11481, SNPs in *KCS4* promoter (rs39451, rs39499, m11502), in *KCS4* genomic region (m11503) and in *KCS4* 3’UTR (m11504). **d**. Tajima’s D value in sliding windows for the genomic region analyzed in **(b)**. Gene models and gene orientation are depicted with grey arrows, *KCS4* is highlighted in yellow. Significantly negative Tajima’s D values are marked with asterisks (*P < 0.05, **P < 0.01).

**Figure 3.**
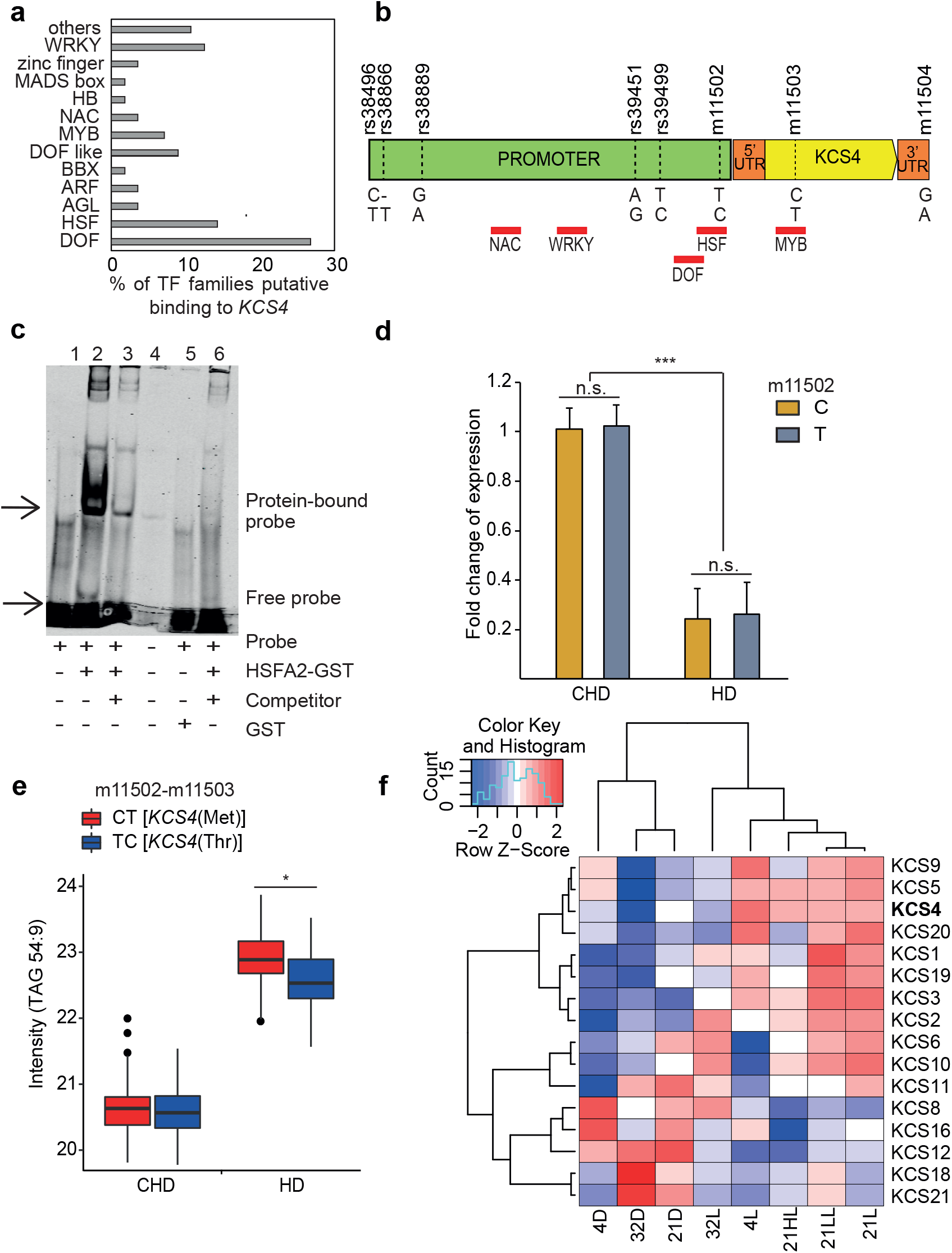
KCS4 transcriptional regulation. **a**. Percentage of transcription factors (TF) classes predicted to bind *KCS4* genomic region according to *in-silico* analysis of binding motives using DAP-seq database (O’Malley et al., 2016). **b**. Position of the binding motifs (red rectangles) for the main TF classes. SNP positions in the promoter and in coding region are marked with horizontal dash-lines and names are stated above gene model. **c**. Electrophoretic Mobility Shift Assay (EMSA) showing the interaction between *HEAT SCHOCK FACTOR A2* (*HSFA2*/AT2G26150) and the HSE binding motif (heat shock element: 5’-nnGnAnnTnCtn -3’) found within the SNP m11502 in *KCS4* promoter region; 1: labelled probe (5⍰-DY682-labelled double-stranded oligonucleotide) only; 2: labelled probe plus HSFA2-GST protein; 3: labelled probe, HSFA2-GST protein and 200× competitor DNA (unlabeled oligonucleotide containing HSE binding site with the “C” allele at m11502), 4: well spill, 5: labelled probe plus free GST, 6: labelled probe, HSFA2-GST protein and 200× competitor DNA (unlabeled oligonucleotide containing HSE binding site with the “T” allele at m11502). **d**. *KCS4* expression for the complete GWAS population in CHD and HD conditions, grouped based on C or T allele at SNP m11502 in *KCS4* promoter region. Expression (fold change) was measured by qRT-PCR. Significant differences between CHD and HD transcript levels and between C and T alleles are shown (*t*-Test: * P < 0.05, ** P < 0.001, *** P < 0.0001, n.s. not significant). Data represent means ± SD (three technical replicates per sample). **e**. Average trait value (TAG 54:9 intensity, in log_2_) for the two major haplotypes determined by SNPs m11502 (promoter) and m11503 (coding region). CT and TC haplotypes encodes for *KCS4*(Met) and *KCS4*(Thr), respectively. Significant differences between haplotypes (*t*-Test: * P < 0.05). **f**. Heatmap of expression pattern for 16 members of the *KCS* multigene family clustered using complete-average method. Transcript abundance was measured 24 hours after treatment. Data was obtained previously (Caldana et al., 2011).

### Dissection of *KCS4* alleles involved in TAG variation, pattern of linkage disequilibrium and signatures of selection

We next mined the full sequence information available for 149 accessions from the GWAS population (http://signal.salk.edu/atg1001/3.0/gebrowser.php), for 80 kb around SNP m11481 on chromosome 1. The sequence alignment (Supplementary File 1) was used to build a SNP matrix (Supplementary Dataset 4) to perform GWAS with more SNPs than those included in the previous experiments using the 250K SNPchip (Horton et al., 2012). We analyzed a total of 1,082 SNPs and short insertion-deletions, in addition to the 131 SNPs previously tested in this region. These analyses aim to discard genetic heterogeneity among other genes in the locus, analyze the pattern of linkage disequilibrium (LD) in the region, determine whether more SNPs in *KCS4* are additionally involved in the variation, and evaluate signatures of selection. The results are shown in Figure 2, exemplified for TAG 54:9. According to the *KCS4* gene model (Figure 2a), we identified 35 additional SNPs in *KCS4* genomic region apart from the 4 SNPs scored in the 250K SNPchip (19 in the promoter, two in the 5’UTR, 13 in the coding region, and one in the 3’UTR; Supplementary Dataset 5). Fewer TAGs with lower LOD values showed association with *KCS4* gene region, as expected when analyzing a reduced number of accessions (< 100) (Supplementary Dataset 5, Figure 2b). The lead SNP using resequencing information was a polymorphism not tested before (rs21933), but in high LD with m11481 (r^2^ =0.73, p << 0.001). In addition, a number of polymorphisms in the upstream region of *KCS4*, not tested in the SNPchip, were identified as being significant, pointing to a putative combined regulation at the promoter region mediated by rs39451, rs39499 and m11502 and at the coding region mediated by the nsSNP m11503 (Figure 2a). Surprisingly, several SNPs in the *KCS4* coding region did not associate (Figure 2b).

All SNPs in the locus harboring KCS4 with high LOD values are in high LD with each other and in high LD with the lead SNPs rs21933 and m11481 (r^2^ > 0.2, p < 0.001, Figure 2c). The remaining SNPs in the *KCS4* coding region are not in LD with the lead SNPs nor with the nsSNP m11503 in the *KCS4* coding region. This result suggests that these mutations may have appeared in a different time during evolution or they may have accumulated later, as a consequence of specific recombination, to increase allele frequency at other SNP positions. To evaluate signatures of selection, we tested Tajima’s D for the 80 kb around m11481 (Figure 2d). Tajima’s D reached significantly negative values along the 80 kb, particularly around SNPs significantly associated in the QTL: rs21933, m11481, rs39451, rs39499, m11502, m11503. This result indicates that the region is under a high selective pressure and supports the selection of *KCS4* as the candidate gene involved in puTAG variation.

### Transcriptional regulation of *KCS4* in Arabidopsis

Several SNPs in the promoter region showed significant LD with the non-synonymous SNP m11503. Therefore, we deepened on the transcriptional regulation of *KCS4* in Arabidopsis (Figure 3). We analyzed 1 kb upstream of the transcription start site for the known transcription factor binding sites using the DAP-Seq database (O’Malley et al., 2016). Several transcription factor (TF) families belonging to land plants could likely bind to the *KCS4* promoter region, indicating that *KCS4* expression is controlled under several regulatory processes (Figure 3a). We furthered our analysis by looking at the core binding sites of the highly enriched TF families that contained binding motifs across SNPs in the promoter region of *KCS4* (rs38889, rs39451, rs39499 and m11502). The top-enriched DOF family (DNA-binding One zinc Finger) does not have a binding site colocalizing with polymorphisms, but in other regions of the promoter (Figure 3b). Similarly, the NAC and WRKY TF families bind to non-polymorphic regions in the *KCS4* promoter, and MYB TF binds near a SNP but in the coding region. Interestingly, HSF (heat shock factor), the second highly enriched TF family has a core binding motif (heat shock element: 5’nnGnAnnTnCtn 3’) within SNP m11502 in the promoter region.

We performed an electrophoretic mobility shift assay (EMSA) to assess the physical interaction of Heat Shock Factor A2 (HSFA2) with the *KCS4* promoter sequence. Additionally, we tested if alleles (C/T) at SNP m11502 influence the binding affinity. HSFA2, a well-studied HSF family member, has been shown to bind to the heat shock element in Arabidopsis (Lämke et al., 2016). As shown in Figure 3c, HSFA2-GST fusion protein bound to the 5⍰-DY682-labeled *KCS4* promoter fragment *in vitro* with the C allele at m11502, resulting in a retardation band on the gel (Figure 3c, lane 2) compared to the free probe (lane 1). Interaction between HSFA2 and the *KCS4* promoter region containing the binding site appears to be specific, as the addition of an excessive unlabeled competitor (same sequence, 200x more) significantly reduced the binding to the labeled probe (Figure 3c, lane 3). Finally, we tested a competitor sequence having the alternative allele at m11502 (T). The latter reduced the binding compared to the labeled probe (Figure 3c, lane 6), suggesting that the T allele at m11502 determines a stronger binding of HSF2A to the *KCS4* promoter region. Overall, this indicates that polymorphisms at the promoter region could exert a dosage effect of transcription factors, influencing the expression of *KCS4*.

Furthermore, we analyzed natural variation in transcript levels of *KCS4* for the CHD and HD conditions in the complete GWAS panel (Figure 3d). *KCS4* transcripts are significantly lower in HD compared to CHD. This correlates significantly negatively with highly unsaturated TAG levels (r^2^ = -0.66, p < 0.001), which increased in HD (e.g., Figure 3e, TAG 54:9). In HD, *KCS4* transcript levels are not significantly different for accessions carrying different alleles at m11502. This confirms that neither differences in TF binding-affinity nor expression levels of *KCS4* determine variation of puTAG levels under HD (Figure 3e). In other words, polymorphisms at *KCS4* promoter region are not the causal for puTAG variation in Arabidopsis natural population.

### Expression patterns of KCS family members under diverse environmental conditions

We analyzed the time-resolved response of Arabidopsis Col-0 accession towards light and temperature for 16 of the 21 gene members using data previously obtained (Caldana et al., 2011). Focusing on 24 h, most *KCS* members, including *KCS4* are upregulated under light conditions and downregulated in darkness, irrespective of the temperature. However, *KCS8, KCS12, KCS16, KCS18*, and *KCS21* showed an opposite behavior. *KCS4* transcript levels are downregulated both under darkness and under HD, but transcript levels are much lower in HD, strengthening the idea of a synergistic effect between light and temperature stress in the *KCS4* response. In addition, *KCS5, KCS9* and *KCS20* showed a similar transcriptional response to *KCS4* across several environmental stimuli (Figure 3f).

### *KCS4* functional validation in puTAG variation

To validate the role of *KCS4* alleles in puTAG variation under HD stress, we selected five accessions: three carrying *KCS4*(Met) and two carrying *KCS4*(Thr). Selection of these accessions was based on puTAG levels measured in the HD experiment (Supplementary Dataset 1). Mutant lines (*kcs4*) were obtained through CRISPR-Cas9 by introducing a 57/58–bp deletion at position +18 from the ATG (Supplementary File 2). This deletion introduced a premature stop codon resulting in a truncated protein of 16 aa compared to the wild-type version of 516 aa. Lines were grown under CHD or HD as described in materials and methods and harvested to perform lipidomic analysis. Since natural variation is detected at a population level, we compared average values for *kcs4*(Met) and *kcs4*(Thr) allelic mutants against wild-type *KCS4*(Met) and *KCS4*(Thr) alleles (Figure 4).

**Figure 4.**
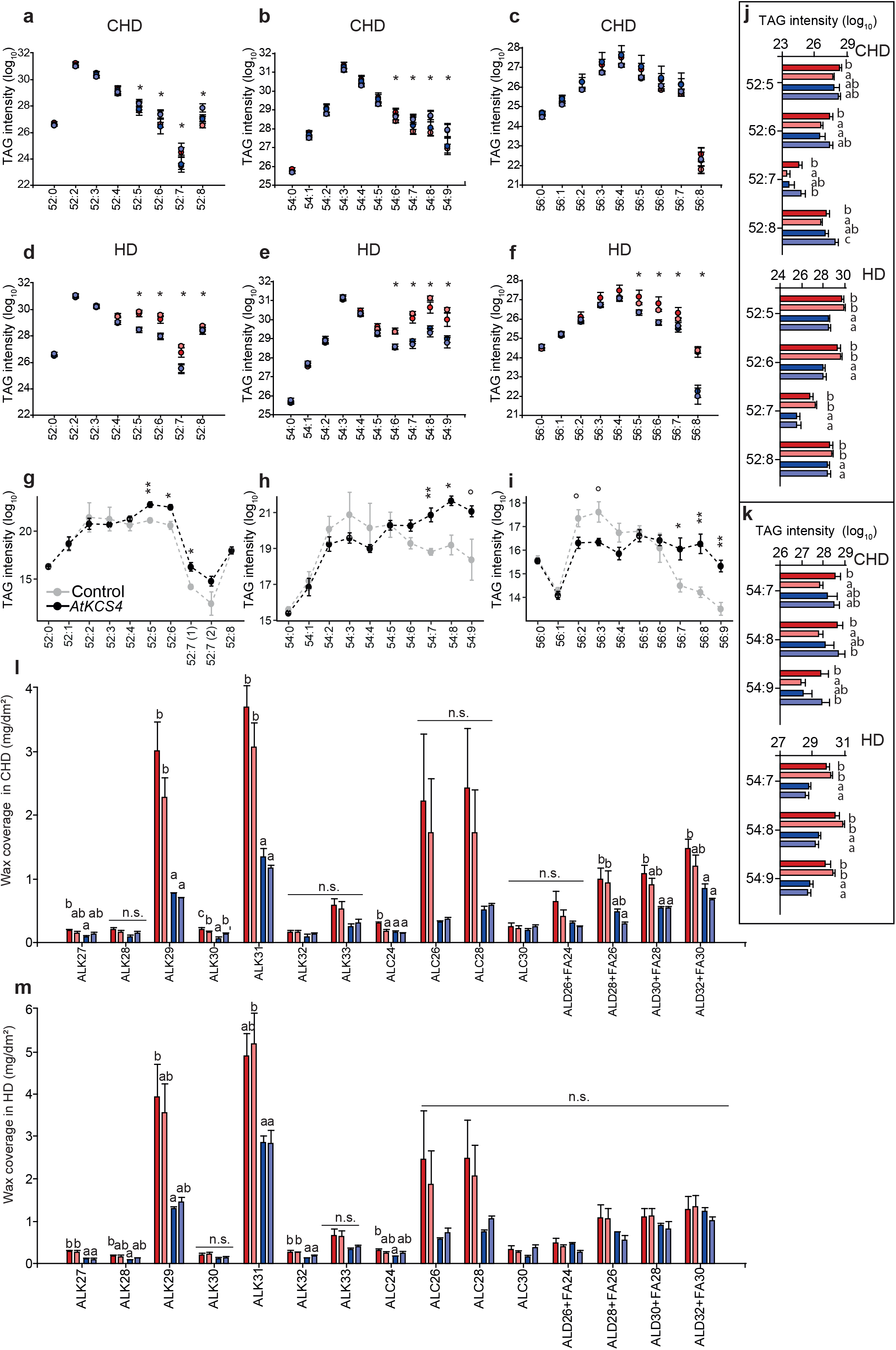
*KCS4* functional validation using allelic mutants in Arabidopsis and heterologous expression in *Nicotiana benthamiana*. **a-f and j-k**. Average TAG profiles for wild-type (WT) accessions and allelic mutants carrying the *KCS4*(Met) allele (WT: red; mutant: pink) and the *KCS4*(Thr) allele (WT: blue; mutant: light-blue) in CHD and HD conditions. One-way ANOVA followed by Tukey post-hoc test was performed to detect differences between lines. Significant changes (* P < 0.05) are marked with asterisks (**a-f**) and with letters for 52 TAG series and 54 TAG series (**j-k**). **g-h-i**. Transient expression of At*KCS4* in *Nicotiana benthamiana* leaves. TAG levels for classes 52 (52:0 - 52:8), 54 (54:0 - 54:9) and 56 (56:0 - 56:0) for the control line (grey circles) and for the At*KCS4* line (black circles). Significant values (*t*-Test) are indicated (° P < 0.1, * P < 0.01, ** P < 0.001). **l-m**. Cuticular wax coverage for the same accessions grown in CHD or HD conditions. One-way ANOVA followed by Tukey post-hoc test was performed to detect differences between lines (* P < 0.05). n.s.: Not significantly different.

Under control conditions (CHD), puTAG levels (X:4 to X:9) are significantly different between *KCS4*(Met) and *kcs4*(Met) lines. Mutants for *kcs4*(Met) presented, in average, lower levels of puTAGs, similar to the levels found for *KCS4*(Thr) accessions (Figure 4 a-c, j-k). This phenomenon is seen for the accessions pooled together as well as for individual accessions (data not shown). On the other hand, puTAG levels did not differ between *KCS4*(Thr) and *kcs4*(Thr). These results support *KCS4*(Met) as the stronger allele and the causal polymorphism for polyunsaturated TAG variation.

Under HD condition, a higher accumulation of puTAGs is seen for accessions carrying *KCS4*(Met) allele (Figure 4 d-f, j-k). However, overall levels of TAGs (saturated and polyunsaturated) increase independently of *KCS4* alleles. This is in agreement with the phenotype observed for the complete mapping panel. Here, mutants don’t differ in puTAG levels from their wild-type counterparts, as if the massive production of puTAGs under stress overrides the solely effect of *KCS4* mutation.

Transient expression of At*KCS4*(Met) in *N. benthamiana* leaves showed significantly higher levels of puTAGs for classes 52, 54, and 56 compared to control plants (Figure 4 g-i). Conversely, TAGs with none or a few unsaturated bonds tend to be higher in control plants. These results show that the transient expression of At*KCS4*(Met) changes the TAG profile in *N. benthamiana*.

In summary, the reduction of puTAGs in the *kcs4*(Met) mutant (Figure 4 a-c, j-k), together with the increasement of puTAGs when *KCS4*(Met) in ectopically expressed in *N. benthamiana* (Figure 4 g-i), confirms that *KCS4*(Met) is the causal polymorphism for polyunsaturated TAG variation and validate GWAS results.

### *KCS4* functional validation in the accumulation of cuticular waxes

VLCFAs are produced by the elongation complex (i.e., KCS, KCR: β-KETOACYL REDUCTASE, HCD: β-HYDROXYACYL-COA DEHYDRATASE and ECR: ENOYL-COA REDUCTASE) in the Endoplasmic Reticulum (ER) and used as precursors for the synthesis of lipid barriers such as cuticular waxes. Therefore, we determined the wax coverage for *KCS4* wild-type alleles and mutants as a proxy of the enzyme activity *in planta* (Figure 4 l-m). Under CHD, lines carrying *KCS4*(Met) showed higher content of waxes than lines carrying *<u>KCS4</u>*(Thr). The mutant *kcs4*(Met) showed a significant reduction of the major components of leaf waxes: C29 and C31 *n-*alkanes, C26 and C28 *n-*alcohols, C30 and C32 aldehydes, C28 and C30 fatty acids. On the contrary, *kcs4*(Thr) does not show differences to its wild-type counterpart. Under HD, the total wax content is increased in the different lines and significant differences are seen between Met and Thr alleles but not between wild type and mutants. These results confirm that the wax coverage and composition is different for both alleles, and that other members of the KCS family cannot fully complement the synthesis of waxes coming from FA elongated by KCS4.

### KCS4 enzymatic activity and cellular localization

*KCS4* shows sequence homology to the remaining 20 members of the Arabidopsis *KCS* multigene family (Joubès et al., 2008). As stated before, KCS enzymes catalyze the first rate-limiting step in the VLCFA elongation, and they determine the substrate specificity.

To evaluate the very-long-chain 3-ketoacyl-CoA synthase activity and substrate specificity of KCS4, we expressed *AtKCS4*(Met) in wild-type yeast and profiled VLCFAs (Figure 5). The strain expressing *AtKCS4*(Met) synthesized significantly higher levels of VLCFA from C20 to C26 compared to the strain transformed with an empty vector (Figure 5a).

**Figure 5.**
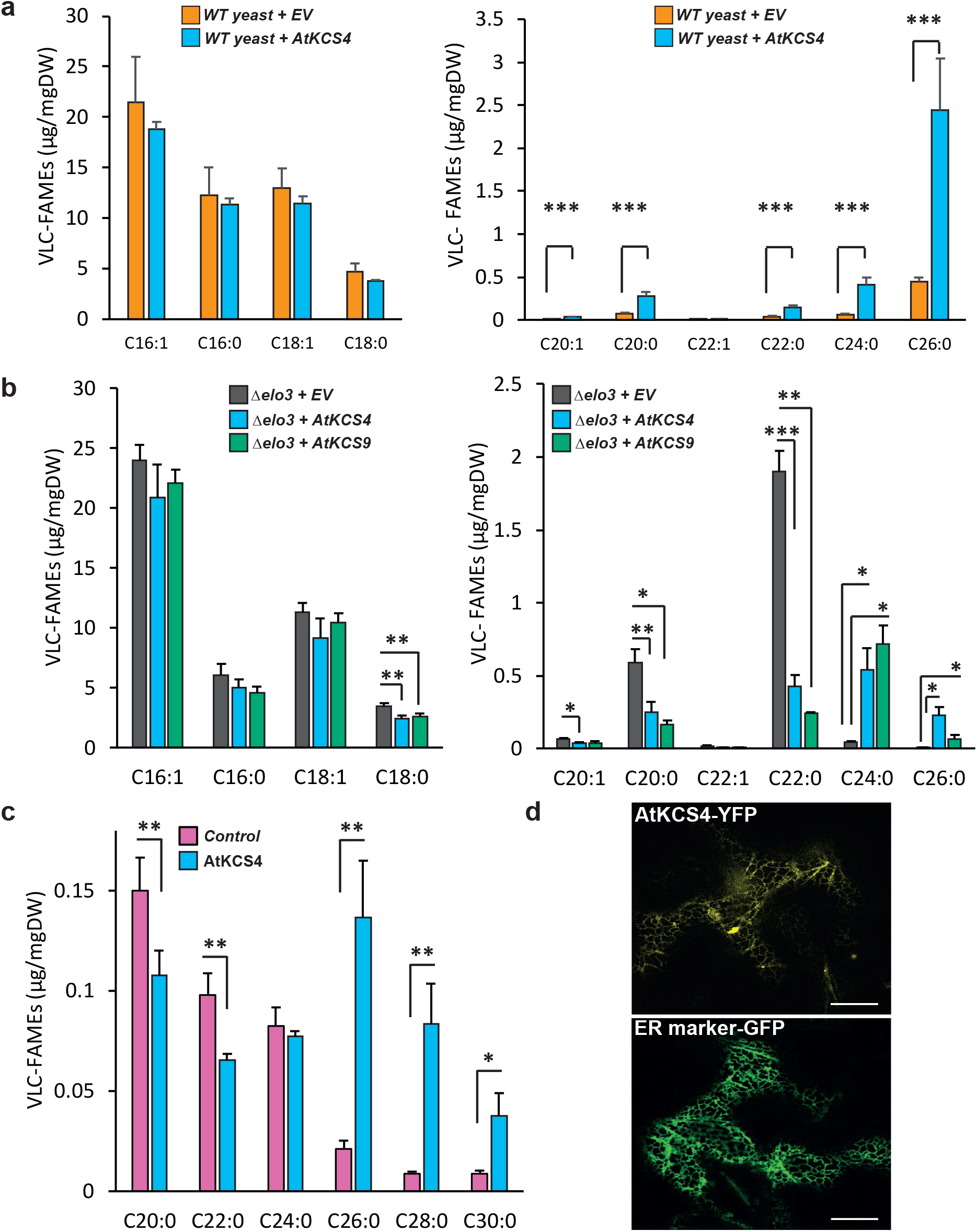
KCS4 activity and specificity in fatty-acid elongation and subcellular localization. **a**. Very long chain fatty acid methyl ester (VLC-FAME) profiles (16 to 18 and 20 to 26, separate panels) of wild-type yeast transformed with an empty vector (orange) and *AtKCS4* (blue). **b**. VLC-FAME profiles (16 to 18 and 20 to 26, separate panels) of Δelo3 yeast mutant transformed with an empty vector (grey), *AtKCS4* (blue bars) and *AtKCS9* (green). **c**. VLC-FAME profiles (20 to 30) from leaves of *Nicotiana benthamiana* control plants (pink) or transformed with *AtKCS4* (blue). Significant differences are shown (*t*-Test: * P < 0.05, ** P < 0.001, *** P < 0.0001). **d**. Confocal images of *N. benthamiana* leaves transiently transformed with KCS4-YFP construct. The plasmid ER-GK CD3-955 was used as an ER marker. Bars = 20 μm.

We further expressed *AtKCS4*(Met) in the Δ*elo3* yeast strain, lacking the endogenous ELO3 elongase (Figure 5b). The Δ*elo3* strain accumulates C20 and C22 VLCFA (Han et al., 2002). Expression of KCS4 led to the elongation of VLCFAs from C22 to C24 and from C24 to C26 (Figure 5b). We also analyzed heterologous expression of At*KCS9* as it is the closest *KCS4* homolog (Figure 5b). KCS9 elongates VLCFAs from C22 to C24 (Kim et al., 2013 and Figure 5b). These results suggest that KCS4 has a differential activity compared to KCS9, pushing further the elongation of VLCFAs up to C26.

Transient expression of *AtKCS4*(Met) in *N. benthamiana* leaves showed a significant decrease of C20 and C22 VLCFAs and a significant increase of VLCFAs from C26 up to C30 compared to the control (Figure 5c). The elongation of VLCFAs up to C30 might be the result of the action of the native elongase complex in *N. benthamiana* due to increased availability of C26 when *KCS4* is transiently expressed. AtKCS4(Met)-YFP was found to label a polygonal and tubular network characteristic of the ER (Figure 5d), in agreement with previous results shown for other KCS family members(Joubès et al., 2008).

We further transformed Δ*elo3* strain with *KCS4* wild-type and mutant alleles (Supplementary Figure 4). Expression of both *KCS4*(Met) and *KCS4*(Thr) alleles led to the elongation of C22 into C24 up to C26 VLCFA in the Δ*elo3* strain with slight differences in the amount of C22 and 2OH-C24 (Supplementary Figure 4a). On the other hand, yeast expressing allelic mutants do not show exactly the composition of the control line, C26 is not accumulated but C22 and C24 VLCFA are still produced at low level compared to yeast expressing wild-type alleles, as if some activity remains. This residual accumulation of VLCFA in the mutants indicates that *kcs4*(Met) and *kcs4*(Thr) lines still have some enzymatic activity, although very weak.

In addition, we performed a time-point experiment at 48, 72 and 96 h for Δ*elo3* strain expressing *KCS4*(Met) *and KCS4*(Thr) wild-type alleles (Supplementary Figure 4b). We could detect that, yeasts expressing *KCS4*(Met) ended up with more C26 and 2OH:C26 than yeasts expressing *KCS4*(Thr), supporting the hypothesis of a differential enzymatic activity between *KCS4* alleles.

KCS1 expression in yeast leads to the production of C18, C20 and C22 VLCFA (Trenkamp et al., 2004; Tresch et al., 2012), KCS2 can elongate from C20 to C22 (Millar et al., 1999; Franke et al., 2009), KCS20 mainly produces C22 and C24 VLCFA, KCS5 and KCS6 mainly produce C24 to C28. All these proteins have different expression profiles under heat and dark stress, some of them upregulated and some of them downregulated (Figure 3f). Expression profiles of KCS4, KCS5, KCS9 and KCS12 are also altered in *kcs4* CRISPR-Cas lines, evidencing a delicate expression balance under stress, between KCS family proteins (Supplementary Figure 5).

### Biomass is affected by KCS4 function

It has been demonstrated that a mild reduction in VLCFA content improve cell proliferation and shoot growth (Nobusawa et al., 2013). To investigate if *KCS4* exert an effect in plant growth, we compared biomass for *KCS4* wild-type accessions and mutant lines. We collected full rosettes from plants grown in CHD and subjected to HD, immediately after treatment. The *kcs4* mutants grew significantly bigger than their WT counterparts (Figure 6). This indicates that biomass is negatively correlated with KCS4 activity and further supports both the idea that mutations at *KCS4* reduce the overall content of VLCFA and that KCS4 activity cannot be overcome by other family members.

**Figure 6.**
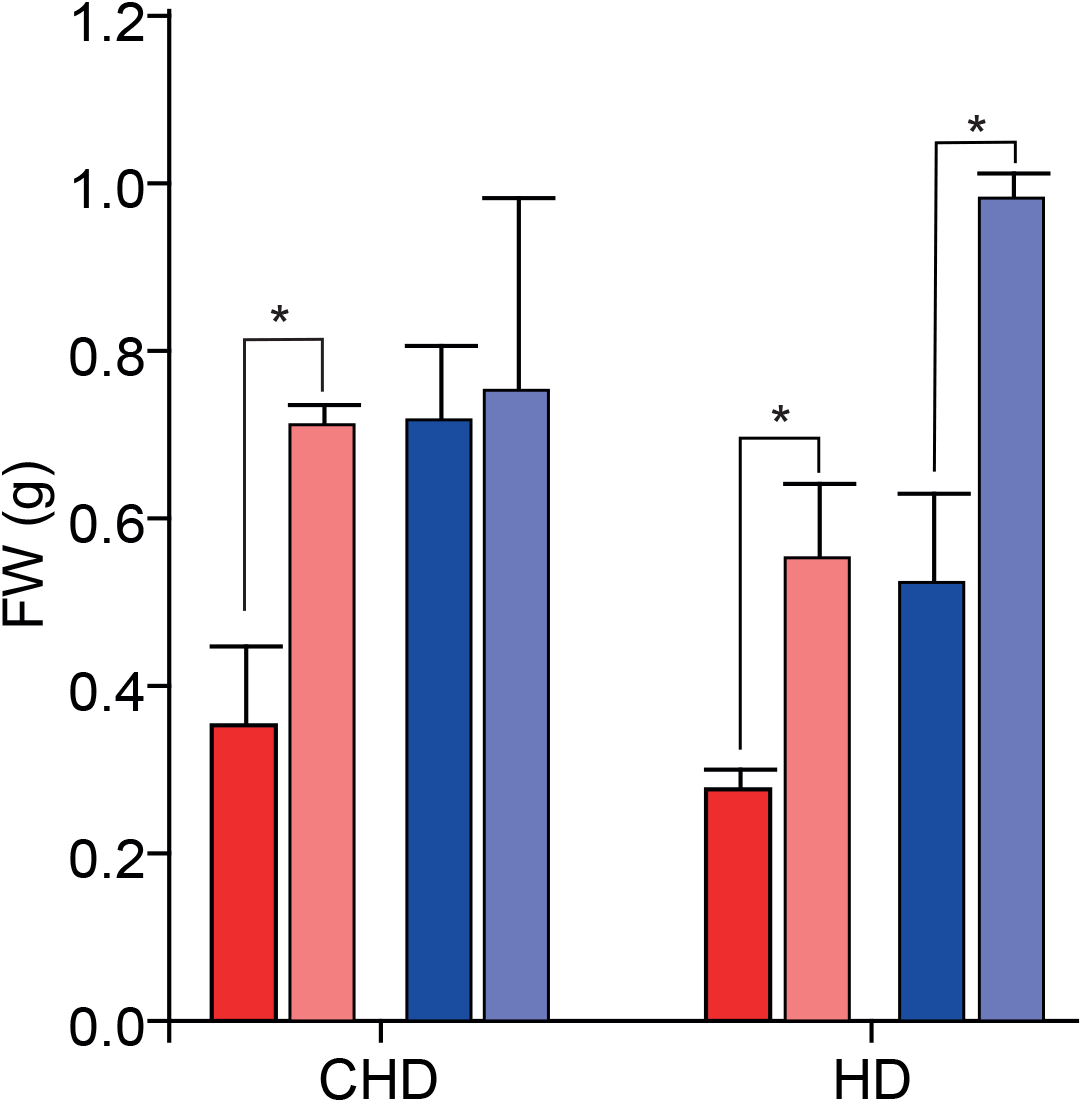
Biomass changes in *KCS4* wild-type and allelic mutants. Rosettes fresh-weight measured in accessions carrying KCS4(Met) or KCS4(Thr), and their respctive allelic mutants. Plants were grown in control conditions (CHD) or subjected to heat and darkness for 24 h (HD) and harvested after the treatment. Significant differences were obtained using *t*-Test (* P < 0.05).

### Phylogenetic signal of the lipidome and correlation with climatic and geographical conditions

To test if there is correlation between lipid profiles and genetic distance (a proxy for phylogeny), K statistics and empirical p-values were calculated for all the measured lipid features in control (CHD) and stress (HD) conditions (Blomberg et al., 2003). The results indicate a correlation between genetic distance and the similarity of lipidomic profiles. Lipids exhibiting a significant phylogenetic signal in CHD are enriched in MGDGs (p ≤ 0.001) while in HD we observed enrichment in TAGs (p ≤ 0.05). Five of the six TAGs with a significant phylogenetic signal are polyunsaturated species associated with the *KCS4* QTL (Supplementary Figure 6).

In addition, climate conditions from the geographical origins of accessions (Fick and Hijmans, 2017) were also analyzed to identify putative correlations with lipidomic profiles. We used a lasso regularized linear model to predict each climate parameter from the lipidomic profiles. In result, we estimated the prediction power for each best-fitting model, and the lipids, being the best predictors for the climate parameters. While the CHD condition did not provide significant results, the HD condition highlighted in total 15 significant model fits, five of which gave also significant results in a cross-validation test (Supplementary Figure 7). All of these five significant variables are related to temperature. In total, six “climate-predictive lipids” were also significantly associated with *KCS4*, with TAG 58:6, TAG 60:6, and TAG 56:8 being linked to the temperature in the driest and the coldest quarter of the year.

**Figure 7.**
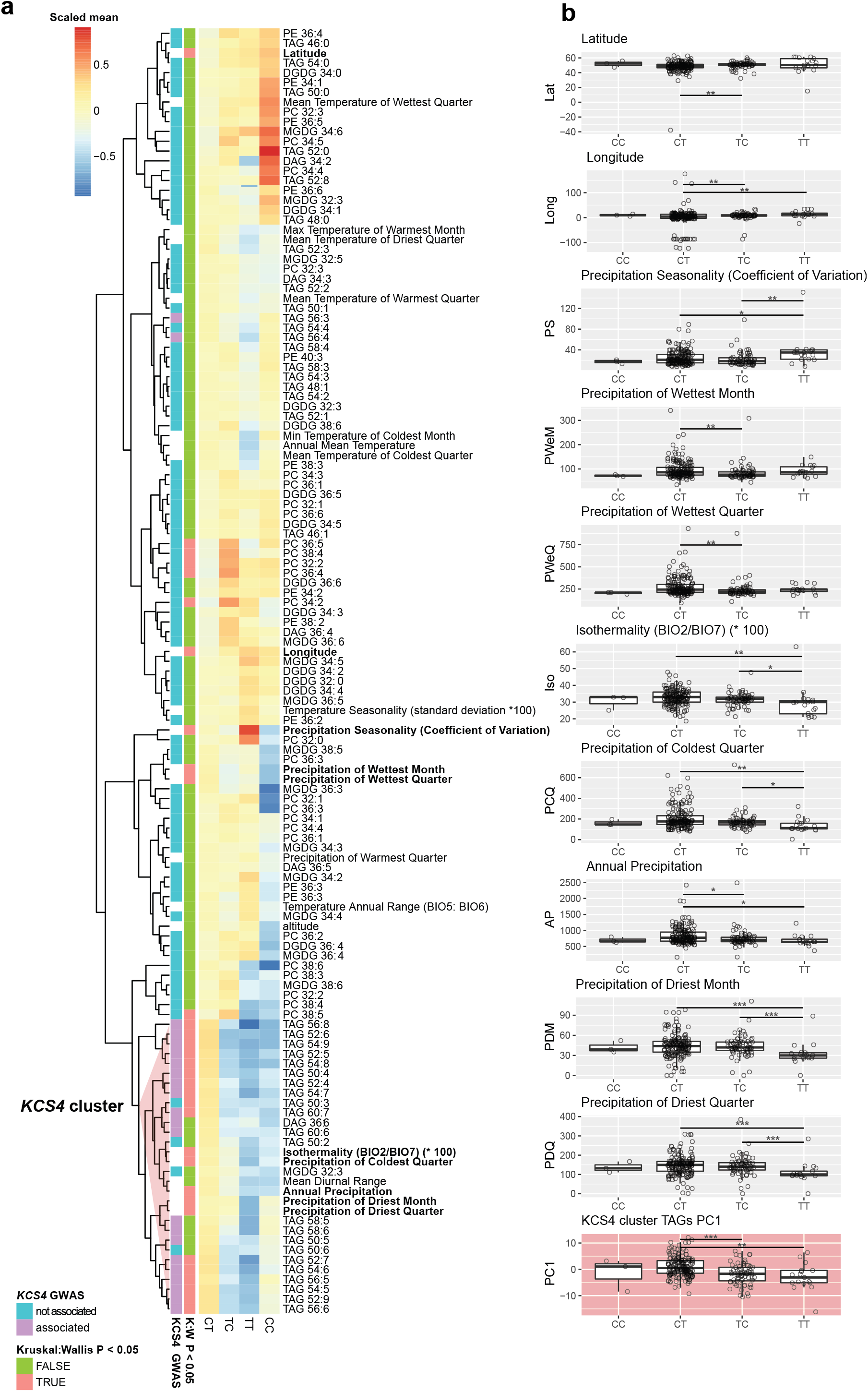
*KCS4* haplotype correlation with climate and geographic parameters. **a**. Heatmap of the mean values of lipid, climate and geographical variables for each of the four KCS4 haplotypes (SNPs m11502-m11503). Results of the *KCS4* locus association as well as nonparametric ANOVA test (Kruskal-Wallis) are highlighted in the row-annotation bar. Cluster dominated by KCS4-associated TAGs is highlighted in red. **b**. Average haplotype value for: the principal component 1 (PC1) obtained using the data in KCS4 cluster (red triangle in **a**), and the climate and geographical parameter showing significant Kruskal-Wallis in **a**. Significant differences are shown (pairwise Wilcoxon rank sum test: * P < 0.05, ** P < 0.01, *** P < 0.001).

Finally, cluster analysis was performed for climate parameters and TAG levels, by grouping accessions according to *KCS4* haplotypes (m11502-m11503: CC, CT, TT, TC), and significant differences between mean values for haplotypes were tested using non-parametric ANOVA (Figure 7). Eight climate parameters plus latitude and longitude differed significantly between haplotypes (p < 0.05, Figure 7a). Moreover, five of them co-clustered with TAGs associated with *KCS4*, four of them related to precipitation and one to temperature (isothermality). The “TT” haplotype in most cases, presented the lowest value for these parameters, suggesting that the allelic effect of these two SNPs is opposite for climate variables (Figure 7b).

## Discussion

Recent years have witnessed the successful application of metabolic GWAS (Luo, 2015). In lipids, the effort has focused more on dissecting the genetic architecture of oil biosynthesis (Correa et al., 2020). Here, we focus on glycerolipid profiles of ∼ 300 Arabidopsis natural accessions grown under control conditions and subjected to a sudden environmental change, given by a combined stress of heat and darkness. Our GWAS exposed the high plasticity of the Arabidopsis genome allowing the identification of genes involved in the regulation of lipid metabolism, in an environment-dependent manner (Figure 1a).

Lipid-based GWAS in control conditions showed several mild-strength associations, similar to what was found for glycerolipids in rice (Hong et al., 2022) and for primary metabolites in maize and Arabidopsis (Wen et al., 2015; Wu et al., 2016), which are controlled by a large number of small-effect loci. This could be explained by the fact that both lipids and primary metabolites are essential for the survival of the cell and thus, do not show on/off phenotypes. This is different from natural variation in secondary metabolites, which are controlled, to a large extent, by major large-effect loci (Chan et al., 2011; Matsuda et al., 2012; Wen et al., 2014; Wu et al., 2018).

In our study, applying combined heat and darkness resulted in a strong perturbation of lipid homeostasis. This led to the identification of a large-effect QTL at Chromosome 1, harboring *KCS4. KCS4* associated with several polyunsaturated TAGs (puTAGs) specifically under carbon starvation conditions (HD, 3D and 6D) in a highly robust manner (Figure 1b, Supplementary Figure 2b, Supplementary Dataset 2/3). Under HD, all puTAGs mapping to *KCS4* are higher than under CHD, but accessions carrying *KCS4*(Met) always have higher levels of puTAGs than accessions carrying *KCS4*(Thr). PuTAGs that mapped to *KCS4* clustered together and correlated negatively to saturated TAGs, further illustrating the role of *KCS4* in the specific regulation of puTAGs (Figure 1c).

We have provided several lines of proof that both *KCS4* is the causal gene for TAG variation and *KCS4*(Met) allele determines higher levels of puTAGs under stresses (HD, 3D, 6D): (i) linkage disequilibrium, correlation among lipid species and haplotypic analysis point *KCS4* as the candidate gene for puTAG variation under stress (Figure 1, Figure 2), (ii) transient overexpression of *KCS4*(Met) in *N. benthamiana* mimics the phenotype we observed in the GWAS population grown under stress conditions (Figure 4 g-i), (iii) lipidomic analysis of allelic mutants (*kcs4*(Met)/*kcs4*(Thr)) and their respective wild-type accessions (*KCS4*(Met)/*KCS4*(Thr)) shows a clear phenotype on puTAG, VLCFA and wax components (Figure 4, Figure 5, Supplementary Figure 5).

KCS4 is a functional enzyme, localized in the ER, with substrate specificity for VLCFAs of C22 and C24 length, and has the ability to elongate VLCFAs to C24 and up to C26 (Figure 5). This result is in agreement with Kim et al. (2021) findings for *kcs4* TDNA mutant, which accumulates C22 and C24. Moreover, the results from ectopic expression in yeast and plants confirmed that KCS4 has specificity for elongation of saturated FAs (Figure 5), in agreement with previous findings that showed that elongase-condensing enzymes are mostly active with saturated FA (Blacklock and Jaworski, 2006).

Under HD, differences in TAG levels for accessions carrying *KCS4* opposite alleles [*KCS4*(Met) vs. *KCS4*(Thr)] become more significant as the number of double bonds in the TAGs increases (Figure 4 d-f, g-i). In addition, TAG levels are significantly different when comparing *KCS4*(Met) with its allelic mutant *kcs4*(Met) in control conditions (CHD) (Figure 4 a-k). In the same line of evidence, the mutant *kcs4*(Met) showed a significant reduction of the major components of leaf waxes compare to its wild-type allele in CHD (Figure 4 l-m). Not unexpectedly, *kcs4*(Thr) did not show any differences in TAG levels of wax compounds compared to *KCS4*(Thr). It has been previously shown in GWAS, that mutating the weaker allele did not always result in clear phenotypic changes for the traits under study (Todesco et al., 2010; Fusari et al., 2017; Wang et al., 2022; Yang et al., 2022).

The phenotype can also depend on the context and a change in either the genetic or the environmental background can amplify or remove the effect (Tonsor et al., 2005). Under HD, there is a strong metabolomic and expression reprogramming as shown before for lipids and primary and secondary metabolites (Burgos et al., 2011; Szymanski et al., 2014; Wu et al., 2016, 2018). This could explain why we cannot observe a phenotype by mutating only one gene. In addition, sometimes effects are not evident when analyzing single mutants, particularly if there are gene families encoding for the same or very similar enzymatic activities. For example, in studies of auxin conjugates, IAA showed a reduction only when triple amidohydrolase mutants were analyzed (Ludwig-Müller, 2011). *KCS4* is one of 21 genes encoding for KCS activity (Joubès et al., 2008; Guo et al., 2016) and we have shown that expression of other KCS members is altered in *kcs4* allelic mutants, indicating a complex regulation between *KCS* genes (Supplementary Figure 5). Of note, the nature of this complex regulation was recently demonstrated to involve protein-protein interactions within the elongase complex (Kim et al., 2022).

During plant evolution, the *KCS* family began with a single gene in algae and evolved via numerous gene duplications into a 21-member gene family in Arabidopsis (Guo et al., 2016). Analysis of LD and Tajima’s D parameters in the region harboring *KCS4* showed that diversity is limited to two major haplotypes and that the region is under strong selective pressure (Figure 2). As *KCS4* is involved in TAG regulation under conditions where carbon reserves are exhausted, it might be possible that *KCS4* has been targeted for natural selection to enhance adaptation in stressful environments. In nature, 24 h of darkness does not occur, but carbon starvation is also observed when plants are exposed to very-limited light periods in regions with high amplitudes of day-length. In agreement with an adaptative role of *KCS4* to environmental stresses, the geographical origin of the accessions (i.e., latitude) correlated with *KCS4* haplotypes (Figure 7). Together, these analyses indicate that polymorphism variation at *KCS4* is important for adaptation in different climatic zones by adjustment of lipid metabolism.

In an event of sudden extended darkness (3D, 6D) or combined heat and darkness (HD), plants become carbon starved (Supplementary Figure 3) (Stitt and Zeeman, 2012). It has been shown that increasing night temperature accelerates enzyme activities exhausting carbon resources prematurely (Pilkington et al., 2015). In this case, FAs are used as alternative substrates for respiration via β-oxidation (Fan et al., 2014). There are many lines of evidence that support the idea that plant lipids are alternative sources of carbon and energy under stress, particularly under carbon starvation. For instance, Kunz et al. (2009) showed that by knocking out the PXA1 transporter, the import of FFA into the perixosome is blocked, resulting in plants dying faster in extended darkness. Fan et al. (2017), through the study of SDP1/LIP1 mutants, could define the crucial role that TAG metabolism has in membrane lipid breakdown, fatty acid turnover, and plant survival under extended darkness. Other study showed that there are different metabolic strategies employed by leaves in response to different darkening treatments (Law et al., 2018). These strategies involve either importing sugars from sink sources, metabolizing amino acids or activating the turnover of lipids from the plastid membranes to the b-oxidation in the peroxisome. The last strategy is particularly important after 6 days of dark treatment. Finally, Yu et al. (2018) demonstrated that the disruption of starch synthesis resulted in a significant increase in FA synthesis via post-translational regulation of the plastidial acetyl-coenzyme A carboxylase and a concurrent increase in the synthesis and turnover of membrane lipids and triacylglycerol.

To summarize our findings, we present a model of lipid remodeling under heat and dark stress in Arabidopsis (Figure 8). In the proposed model, darkness induces the predominant response within the combined stress, blocking *de novo* FA synthesis (*de novo* FAS) and producing considerable membrane-lipid remodeling. We observed decreased levels of MGDGs and DGDGs across accessions, as previously shown (Burgos et al., 2011). Under carbon starvation, FAs, mostly 16:3 and 18:3, are released from galactolipids by lipases (Higashi et al., 2015; Fan et al., 2017). FAs are then exported to the ER, where KCS4 is localized and acts as a branch point in the fate of FAs, directing the saturated ones to the very-long-chain fatty-acid pathway (VLCFA). Two *KCS4* alleles have differential capacity to allocate saturated FAs into the VLCFA pathway, with a major -more efficient-allele, *KCS4*(Met), and a minor -weaker-allele, *KCS4*(Thr). *KCS4*(Met) accessions are, as well, efficiently channeling the saturated FA from the Acyl-CoA pool to cuticular waxes, as they present higher accumulation of waxes than *KCS4*(Thr) accessions under stress (Figure 4). Polyunsaturated TAG levels are, therefore, higher in *KCS4*(Met) accessions comparing to *KCS4*(Thr) accessions, as the result of a rise in the ratio of polyunsaturated-to-saturated FAs that proceed to TAG formation (Figure 4). This effect of *KCS4*(Met) on TAG levels is only observed under stress conditions, when there is an increased amount of free-polyunsaturated fatty acids coming from membrane lipids, and when it necessary to sequestered them into TAGs, to protect the cell from the free-fatty-acid-toxicity induced damage (Kao et al., 2018).

**Figure 8.**
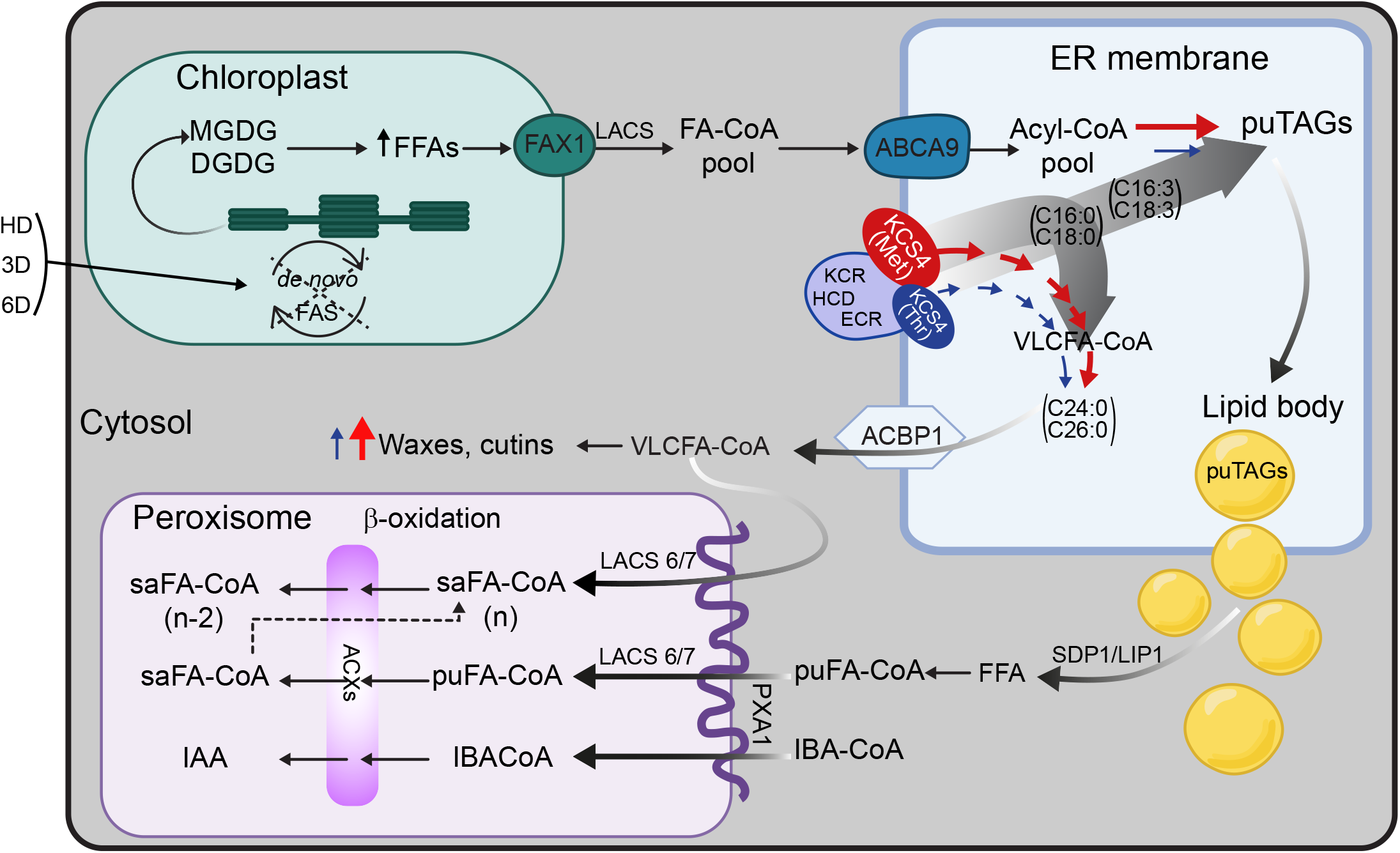
Model summarizing the role of KCS4 on FA fate in Arabidopsis under carbon starvation. The lack of de novo FA synthesis under HD induces the degradation of galactolipids (MGDGs and DGDGs) from the chloroplasts’ thylakoids. FFAs are then exported to the ER (FA-CoA pool), where KCS4 is localized and acts as a branch point in the fate of FAs, channeling saturated FAs to VLCFA elongation and tipping the balance to a higher polyunsaturated-to-saturated-FA ratio for puTAG accumulation (degrade grey arrows). Two *KCS4* alleles have differential capacity to allocate saturated FAs into the VLCFA pathway, with a major -more efficient-allele, *KCS4*(Met), and a minor - weaker allele, *KCS4*(Thr), exemplified here as red and blue and in different sizes, respectively. *KCS4*(Met) accessions efficiently channel the saturated FA from the Acyl-CoA pool to cuticular waxes, as they present higher accumulation of waxes than *KCS4*(Thr) accessions under stress. In addition, polyunsaturated TAG (puTAGs) levels are, therefore, higher in *KCS4*(Met) accessions comparing to *KCS4*(Thr) accessions, as the result of a rise in the ratio of polyunsaturated-to-saturated FAs going to TAG formation. VLCFA-CoA produced by the elongase complex (here exemplified by KCS4, KCR, HCD, ECR) go to increase the cuticular waxes under HD, and/or directly serve as a source of carbon undergoing degradation in the peroxisome; whereas puTAGs accumulated into lipid bodies first. FFAs are then released from lipid bodies by the action of lipases (SDP1/LIP1) and further imported to the peroxisome to undergo β-oxidation. Acronyms: ABCA9 (ATP-BINDING CASSETTE A9), ACBP-1 (Acyl-CoA-binding protein 1); ACX (ACYL-COA OXIDASE); CoA (Coenzyme A); DGDG (Digalactosyldiacylglycerol); ECR (ENOYL-COA REDUCTASE); FAs (fatty acids); FAS (fatty-acid synthesis); FAX1 (FATTY ACID EXPORT1); FFAs (free fatty acids); HCD (β-HYDROXYACYL-COA DEHYDRATASE); IAA (indol-3-acetic-acid), IBA (indol-3-butyric-acid), KCR (β-KETOACYL REDUCTASE); KCS4(Met/Thr) (3-KETOACYL-COA SYNTHASE4); LACS (LONG-CHAIN ACYL-COA SYNTHETASE); SDP1/LIP1 (LIPASE 1); MGDG (Monogalactosyldiacylglycerol); PXA1 (PEROXISOMAL ABC-TRANSPORTER 1); pu (polyunsaturated), sa (saturated), TAGs (Triacylglycerols); VLCFA (Very-Long-Chain-Fatty-Acid).

Our work demonstrates that KCS4 is a key enzyme involved in the regulation of puTAG levels under heat and darkness and under extended darkness alone, adding complexity to lipid-metabolism regulation under carbon starvation. The enzymatic function of KCS4 in the synthesis of VLCFA is affecting the accumulation of the unsaturated Acyl-CoA pool (16:3 and 18:3, for example) and the sequestration of them into puTAGs.

In conclusion, our study demonstrates that lipidomic-GWAS performed under stress conditions is a powerful and unbiased tool for shedding light on the genetic control of lipid metabolism. We identified a QTL to which lipids mapped in an environment-specific fashion, thus illustrating the plasticity of the genome in the regulation of lipid metabolism in changing environments. This approach fleshed out the fundamental role that KCS4 has in FA channeling under stress and proved to be powerful in unraveling the genetic regulation of complex phenotypes.

## Methods

### Plant material

The GWA mapping population comprised a total of 330 *Arabidopsis thaliana* natural accessions belonging to the HapMap panel (Horton et al., 2012). This panel was selected because the accessions maximize diversity and minimize redundancy and close family relatedness (Baxter et al., 2010).

### Experimental design

Accessions or mutant lines were grown in soil (potting compost) in short days (8 h light) for four weeks, then transferred to long days (16 h light) for an additional period of two weeks. Control plants were harvested immediately and stressed plants were subjected to the treatment and harvested right after it.

In control conditions (for heat/dark experiment: CHD, for dark experiment: CD), plants were grown in long days (16h light, 8 h dark), temperature was kept at 21/16°C (day and night, respectively), light intensity at 150 μE m^-2^ sec^-1^, humidity 75%. In the heat and dark experiment (HD), plants were grown like the plants in CHD for six weeks, then subjected to dark and 32°C continuously for 24 h, and then harvested. In the dark treatment, plants were grown like the plants in CD for six weeks and then transferred to continuous darkness either for 72 h (3D) or 144 h (6D). Plants from HD, 3D, 6D were harvested 24 h, 72 h, and 144 h after the beginning of the treatment, respectively. Between three and six plants from each accession/line were collected for each condition tested.

### Targeted lipid profiling by LC–MS and data acquisition

Lipid extraction and measurement from Arabidopsis leaves (100 mg for each sample) were performed as described previously (Hummel et al., 2011). In brief, dried organic phase was measured using Waters Acquity ultra-performance liquid chromatography system (Waters, http://www.waters.com) coupled to Fourier transform mass spectrometry (UPLC–FT–MS) in positive ionization mode. Analysis and processing of raw MS data were done with REFINER MS® 10.0 (GeneData, http://www.genedata.com). Workflow included peak detection, retention time alignment and removal of chemical noise and isotopic peaks from the MS data. Obtained mass features characterized by specific peak ID, retention time, m/z values and intensity were further processed using custom scripts in R (R Core Team, 2019). Clusters with mean of signal intensities lower than 50000 were removed and only peaks present in at least 70% of the samples were kept for further analysis. Peak intensities were normalized by the day of measurement and median and subsequently log_2_ transformed. After that, obtained molecular features were queried against a lipid database for annotation. The in-house database used includes 219 lipid species of the following classes: diacylglycerols (DAGs), digalactosyldiacylglycerols (DGDGs), monogalactosyldiacylglycerols (MGDGs), phosphatidylcholines (PCs), phosphatidylethanolamines (PEs) and triacylglycerols (TAGs). This was carried out by comparing their retention times and exact masses against those of the reference compounds, allowing maximal deviations of 0.1 minutes and 10 ppm. Identified lipids were confirmed by manual verification of the chromatograms using XCalibur (Version 3.0, Thermo-Fisher, Bremen, Germany) (Alseekh et al., 2021).

### General statistical and multivariate analysis of lipidomic data

To calculate the correlation between lipids, Pearson correlation was used. The correlation plots and heatmaps were created using the package pheatmap (Raivo Kolde, 2019). The lipids were clustered using complete-linkage clustering. Principal component analysis (PCA) was obtained from the correlation matrix of lipidomic data (CHD and HD together) using the pcaMethod (Stacklies et al., 2007) package.

To detect the fold change in lipid-feature levels across all accessions for each lipid feature, fold change was calculated by dividing the average of the five highest intensities measured in a given accession by the average of the five lowest intensities, A) for all annotated lipids (N_CHD_ = 98, N_HD_ = 109), B) only for TAGs (N_CHD_ = 30, N_HD_ = 41), C) for lipid classes other than TAGs (N_CHD_ = 68, N_HD_ = 68).

### GWAS analysis

Data acquisition for GWAS, mapping and locus identification were performed as described previously (Wu et al., 2016; Fusari et al., 2017). Genome-wide association analysis (GWAS) was performed using a compressed mixed linear model (MLM) implemented in the GAPIT (Lipka et al., 2012) package. The 109 (HD), 98 (CHD), 80 (3D), 109 (6D), and 122 (CD) lipid features were used as phenotypes (Supplementary Dataset 1), whereas a set of 214,051 SNPs was used as genotypic data (250K SNPchip). A total of three principal components were included in the MLM to account for population structure and the SNP fraction considered to perform PCA was set to 0.1 to avoid excessive computations. Kinship matrix and other parameters were set to default values.

Additionally, we performed GWAS using full genome sequence from a section of chromosome 1. Shortly, sequences for 80 kb around *KCS4* (AT1G19440, TAIR10 position: 6689119 - 6769118) were obtained from the Arabidopsis 1001 Genomes project (http://signal.salk.edu/atg1001/3.0/gebrowser.php, 162 accessions). The alignment was performed using the default routine implemented in MAFFT (Katoh et al., 2019) with a final manual inspection to discard alignment problems due to big gaps or missing data (Supplementary file 1). For nine accessions, two sequences were available. The sequences included further in the analysis were selected based on agreement of alleles with the SNPchip data and/or on % of non-missing data (≥ 90%). Final SNP matrix, obtained by filtering for minor allele frequency ≥ 5%, contained 1,213 SNPs and indels (Supplementary Dataset 4). This matrix was used as the genotypic data. Population structure and cryptic relationships among accessions were taken into account by including principal components as cofactors and a kinship matrix obtained with SNPs in the 250K SNPchip for the subset of 162 accessions (Supplementary Dataset 4). Lipid features collected formerly (HD and 3D) Supplementary Dataset 1) were used as phenotypes. The three data sets (genotype, population structure and phenotypes) matched for 149 accessions in the HD experiment and for 134 accessions in the 3D experiment. MLM was implemented using TASSEL software with the default settings for compressed MLM (Bradbury et al., 2007).

### SNP filtering, QTL identification and candidate gene selection

For results obtained using GAPIT package, SNPs associated with lipid species having a p value < 1/N (N=214,051) were extracted. SNPs were assigned to the same QTL if the genomic distance between them was ≤ 10 kb. All genes in a given QTL were taken as putative candidates. Further candidate gene selection for validation was done based on functional annotation or based on previous knowledge.

For results obtained using Tassel software, SNPs were regarded as significantly associated with lipid traits after setting Bonferroni threshold (α = 0.05/N, N = 1,213 polymorphism tested).

### Linkage disequilibrium, nucleotide diversity, and haplotype analysis

Linkage disequilibrium (LD) was computed as the coefficient of correlation between pair of sites (r^2^) using the LDheatmap package (Shin et al., 2006), either with the SNPchip (for 1,300 and 314 accessions) or with the full sequence data (for 149 accessions). For deviations from neutrality, Tajima’s D statistic (Tajima, 1989) was computed as described previously (Fusari et al., 2017).

Haplotypes for *KCS4* were obtained by using SNPchip data considering 1,000 bp upstream the ATG, the coding region and 1,000 bp downstream the stop codon (SNPs= m11502, m11503, m11504, m11505). Accessions belonging to the same haplotype were grouped and the haplotype median was obtained for each lipid feature. To cluster haplotypes, a combination of allele sharing distance and Ward’s minimum variance was used, as described before (Gao and Starmer, 2007; Qiu et al., 2015). One-way ANOVA followed by Bonferroni correction for multiple comparisons was performed to test differences between haplotypes (p < 0.05).

### Mutant line selection and genotyping

For Arabidopsis CRISPR-Cas9 lines, the binary vector containing multiple gRNAs and the Cas9 nuclease were generated based on a classic cloning procedure described previously (Schumacher et al., 2017). A pair of guideRNAs was used to target the N-terminal coding sequence of *KCS4* using the spacer sequences indicated in Supplementary Dataset 8. As background, wild-type accessions carrying *KCS4*(Met): Pro-0, Petergof, Per-1, Col-0; and *KCS4*(Thr): Si-0, Bu-8, Sh-0 were chosen based on TAG levels phenotyped in HD. Mutations were confirmed by PCR and sequencing of *KCS4* using primers listed in Supplementary Dataset 7. Sequence contigs and alignment were done using Bioedit (Hall, 1999).

For heterologous expression in yeast and transient expression in *Nicotiana benthamiana, KCS4* and *KCS9* open reading frame was amplified from Arabidopsis flower cDNA using primers listed in Supplementary Dataset 7. The corresponding PCR fragments were inserted into the pDONR™221 ENTRY vector using the GATEWAY® recombinational cloning technology (Invitrogen, Darmstadt, Germany) and subsequently transferred into pK7YWG2, pK7WG2D and pvtLEU DESTINATION vectors (Karimi et al., 2002; Dittrich-Domergue et al., 2014) by using LR clonase II. All transient heterologous expressions in *N. benthamiana* were made using the *Agrobacterium tumefaciens* strain GV3101 and five-week-old plants cultivated in controlled conditions (16 h light photoperiod, 25 °C) (Perraki et al., 2012). *N. benthamiana* plants were agroinfiltrated on the abaxial side of the leaves and analyzed 2 and 5 days after agroinfiltration for subcellular localization experiments and for overexpression, respectively.

For yeast experiments, *Saccharomyces cerevisiae* strains INVSc1 [MATa *his3Δ1 leu2 trp1-289 ura3-52*] and Δ*elo3* [MATa *his3Δ1 leu2 trp1-289 ura3-52 elo3Δ*] were used to express *KCS4* wild-type and allelic mutants and *KCS9*.

### qRT-PCR and eQTL analysis

*RGS1-HXK1 INTERACTING PROTEIN1/RHIP1* (AT4G26410) was used as reference gene for *KCS4* transcript analysis in CHD and HD for the complete GWAS population (Czechowski et al., 2005). *GAPDH3* (AT1G13440) was used as reference gene for starvation markers and *KCS* transcripts analysis. All primers used are listed in Supplementary Dataset 7.

qRT-PCR was performed on three biological replicates per ecotype. Each sample was tested in three technical replicates. qRT-PCR was performed on an ABI Prism® 7900 HT real-time PCR system (Applied Biosystems/Life Technologies, Darmstadt, Germany) in 384-well PCR plates with a total reaction volume of 5μL (2 μL forward and reverse primer mixture, 0.5 μM each), 0.5 μL cDNA and 2.5 μL Power SYBR® Green-PCR Master Mix (Applied Biosystems/Life Technologies) using the following cycling program: 50°C for 2 min, 95°C for 5 min, 40 cycles of 95°C for 15 s and 60°C for 60 s, followed by a denaturation step of 95°C for 15 s, 60°C for 15 s, followed by a continuous temperature increase (0.3°C per s) to 95°C (for 15 s). Steps 5, 6 and 7 were introduced to record a dissociation or melting curve for each product in order to detect non-specific amplification. Expression values were calculated by 2^(-ΔΔCt)^ using either *ACTIN2* or *GAPDH3* as reference genes. Errors are given as the lower (2^-(ΔΔCt-SD)^) and upper (2^-(ΔΔCt+SD)^) limit of the standard deviation around the mean.

### EMSA

EMSA reactions were performed as previously described (Thirumalaikumar et al., 2018). Briefly, HSFA2-GST proteins were prepared as stated before (Puranik et al., 2011). *HSFA2* coding sequence was PCR-amplified from Arabidopsis heat-stressed cDNA using primers listed in Supplementary Dataset 7. PCR products were recombined in pDEST24 using the GATEWAY cloning system (Invitrogen). Recombinant vectors were transformed into *Escherichia coli* Star (DE3) pRARE generated by transforming the pRARE plasmid isolated from Rosetta (DE3). 5’-labeled with fluorescent dye DY682 and unlabeled *KCS4* promoter regions spanning the SNP position m11502 (C or T allele) were purchased from Eurofins MWG operon. DNA EMSA reactions were performed using Odyssey Infrared EMSA kit (Li-COR, Bad Homburg, Germany). Competitor was added at 200x the amount of probe.

### Fatty-acid composition and content analysis for *N. benthamiana and yeast*

For fatty acid methyl esters (FAMEs) analysis in *N. benthamiana*, three agroinfiltrated leaf discs were transferred to tubes containing 1 mL 0.5 M sulfuric acid in methanol containing 2% (v/v) dimethoxypropane and 20 μg of heptadecanoic acid (C17:0) as internal standard. Transmethylation was performed at 85°C for 3 hours. For FAMEs analysis in yeast, yeasts were grown in 5 mL of appropriate liquid minimal medium for 1 week at 30°C, then pelleted and washed in 2 mL of 2.5% (w/v) NaCl. Yeast cell pellets were resuspended in 1 mL 0.5 M sulfuric acid in methanol containing 2% (v/v) dimethoxypropane and 20 μg of heptadecanoic acid (C17:0) as internal standard and transmethylation was performed at 85°C for 1 hour. After cooling, 2 mL of 2.5% (w/v) NaCl was added, and fatty acyl chains were extracted in 300 μL of hexane. Samples were analyzed by GC-FID and GC-MS as previously described (Domergue et al., 2010).

### Cuticular wax analysis

Epicuticular waxes were extracted from leaves by immersing tissues for 30 s in chloroform containing docosane (C22 alkane) as internal standard. Extracts were derivatized and analyzed as previously described (Bourdenx et al., 2011).

### Confocal microscopy

Live imaging was performed using a Leica SP5 confocal laser scanning microscopy system (Leica, Wetzlar, Germany) equipped with Argon, DPSS, He-Ne lasers, hybrid detectors and 63x oil-immersion objective. Two days post agroinfiltration, *N. benthamiana* leaf samples were gently transferred between a glass slide and a cover slip in a drop of water. The plasmid ER-gk CD3-955 was used as a fluorescent ER marker (Nelson et al., 2007). YFP and GFP fluorescence were observed using excitation wavelength of 488 nm and their fluorescence emission was collected at 490–540 nm.

### Biomass determination

Plants were grown and treated as described for lipid measurements. Plants from wild-type accessions carrying *KCS4*(Met) and *KCS4*(Thr) as well as allelic mutants *kcs4*(Met) and *kcs4*(Thr) were grown in CHD or treated for 24 h with HD and harvested right after (CHD and HD at the same time). Biomass measurement was performed on complete rosettes (n=3 ± SD) for each line. Results from accessions or mutants were then pooled according to the allele carried.

### Geographic and phylogenetic signal of the lipidome

The neighbor-joining tree was constructed by BioNJ algorithm (Gascuel, 1997), using J-C distance in SeaView 4.4.3 software (Gouy et al., 2010). Phylogenetic signal was computed using K statistics and as a background a Brownian motion model of trait evolution (Blomberg et al., 2003) in the ‘picante’ R package (Kembel et al., 2010). Significance of the phylogenetic signal was estimated by comparison of observed variance patterns to a null model obtained by 9,999 random permutations of the accession labels across the phylogeny. The altitude at the sites was estimated using the geomapdata database (Lees, 2000) based on the latitude and longitude coordinates. The climate conditions have been downloaded from the WorldClim database (www.worldclim.org (Fick and Hijmans, 2017)). The lipidomic data were regressed against the geodesic distance matrix using non-parametric multivariate variance analysis (Anderson, 2001). The relationship between lipid levels and the climate parameters was analyzed using regularized regression model (Friedman et al., 2010). The regularization parameter alpha and penalty parameters lambda were estimated using 10-fold cross-validation within the training set. In result the Elastic Net regularization with parameter alpha = 0.5 was selected as optimal and applied for all regression models and the ‘λmin + 1SE’-rule was applied to choose the penalty parameter for each model separately. Prediction power was estimated in Monte Carlo cross-validation test with 100 samples with 57% and 25% of samples taken as a training and test set respectively. Kruskal-Wallis and Wilcoxon rank sum tests have been performed using the R stats package.

## Author Contribution

YB and CMF conceived the project and supervised the study. UL conducted GWAS experiments, obtained and selected KO mutants, performed experimental validation with KO, statistical analysis and interpretation of results. A-KR help UL in experimental work and performed analysis of the phenotypic data. JJ and MB conducted experiments on tobacco and yeast lines. JSz conducted the phylogenetic and geographical-signal analysis. VPTK provided expertise and assistance on TF analysis and advised on EMSA experiment and qRT-PCR assay. ML and SW assist in GWAS and multivariate analysis. FZ provided the plant material and carried out the lipid profiling for 3D and 6D GWAS. NE, SK and SM helped UL in qRT-PCR assay for GWAS population and KO validation. HX and WJ performed lipid profiling for *kcs4* mutants. JSc obtained KO constructs for transformation of Arabidopsis. AS provided access to LC-MS. ARF and AS help with discussion. CMF performed TASSEL, LD, Tajima’s D and haplotype analyses, analysed and interpreted the results and helped with data visualization. CMF, UL and YB wrote the paper with input from, JJ, JSz and YL-B. All authors read and draft the manuscript.

## Acknowledgements

We would like to thank Prof. Dr. Lothar Willmitzer for discussion, support and the opportunity to develop the project. In addition, we acknowledge Änne Michaelis and Gudrun Wolter for their outstanding technical assistance in LC-MS measurements, and Dr. Karin Kohl for her help in plant cultivation. We thank Hezi Tenenboim for his editorial aid. CMF is a career member of the National Research Council (CONICET) and leader of a Max Planck Partner Group. This research was supported by the Israel Science Foundation (grant no. 859/19).

## Figure legends

**Supplementary Figure 1. Phenotypic analyses for the Arabidopsis GWAS panel under control and stress conditions**

**a**. Principal component analysis (PCA) of lipid data for control condition (CHD, blue circles) and heat and dark condition (HD, red diamonds). Each dot represents a single accession. PC1 and PC2 explain together 67.5% of the variance observed in the phenotypic data. **b**. Heatmap of differential accumulation of lipid levels in stress (HD) and control (CHD) conditions. Lipids are clustered by lipid class. For each accession, the lipid level in HD was subtracted by the lipid level in CHD. Positive values (red) denote higher levels of lipid in HD condition, negative values (blue) denote higher level of lipid in CHD condition (scale= log2 (Δ of lipid intensity). **c**. Correlation pattern among lipid levels in both conditions: CHD (blue rectangle) and HD (red rectangle). Pairwise Pearson correlations (r^2^) were calculated between each lipid across all accessions. Lipids are highlighted with colors according to lipid classes (references included in the figure). **d**. Distribution of fold change of lipid⍰feature levels across all accessions. For each lipid feature, fold change was calculated by dividing the average of the five highest intensities measured in a given 5 accessions by the average of the five lowest intensities. Y-axes is expressed in percentage of (A) all metabolite features (N=109 in stress, N=98 in CHD), (B) only TAG features (N=41 in stress, N= 30 in CHD) and (C) other lipid features (N=68 in stress and N=68 in CHD).

**Supplementary Figure 2. Results from GWAS analysis on lipidomic data from plants grown under extended darkness (3D, 6D) conditions**.

**a**. Chromosome scheme and Quantitative Trait Loci (QTL) identified in the two experimental conditions (3D = circles, 6D = squares), for different lipid classes: phosphatidylethanolamine (PE), phosphatidylcholine (lecithin) (PC), monogalactosyldiacylglycerol (MGDG), digalactosyldiacylglycerol (DGDG), diacylglycerol (DAG) and triacylglycerol (TAG). Color/shape references are included in the figure. Gene IDs are included only for QTL co-localizing for two or more lipid species with LOD score > 5.3. **b**. Manhattan plots obtained for TAG 54:9 in 3D (left) and in 6D (right). **c**. Average trait value (intensity of TAG 54:9, log_2_ scale) for the different *KCS4* haplotypes using SNPs m11502, m11503, m11504 and m11505. **e**. TAG 54:9 average value for accessions carrying the KCS4(Met) and KCS4(Thr) alleles.

**Supplementary Figure 3. Quantification of starvation markers in KCS4 wild-type accessions and allelic mutants**.

Transcripts were measured in 5 accessions, 3 carrying *KCS4*(Met) and two carrying *KCS4*(Thr), and five allelic mutants from those 5 accessions. Plants were grown in control conditions (CHD) or subjected to heat and darkness for 24 h (HD) and harvested both group at the same time point. Three biological replicates were used for each line, results from separate accessions were pooled according to the allele carried. Fold change of expression was obtained using the 2^-(ΔΔCT)^ method using GAPDH3 (At1g13440) as housekeeping gene. Statistical significance was performed with ANOVA followed by Fisher LSD. Means with a common letter are not significantly different (P > 0.05).

**Supplementary Figure 4. KCS4 enzymatic activity**

**a**. Very long chain fatty acid methyl ester (VLC-FAME) profiles (22 to 26) of Δelo3 yeast mutant transformed with wild-type At*KCS4*(Met)/At*KCS4*(Thr) and allelic mutans *kcs4*(Met)/*kcs4*(Thr). References for colors are in the figure. Significant differences between alleles obtained with *t*-Test: ° P < 0.1, ** P < 0.01, *** P < 0.001. **b**. VLC-FAME profiles (22 to 26) at different times of harves (48, 72 and 96 h) for At*KCS4*(Met) and At*KCS4*(Thr) wild-type alleles. Significant differences were obtained with ANOVA followed by Tukey test (P < 0.05).

**Supplementary Figure 5. Quantification of *KCS4, KCS5, KCS9* and *KCS12* transcripts in wild-type *KCS4*(Met)/*KCS4*(Thr) accessions and kcs4(Met)/kcs4(Thr) allelic mutants**.

Transcripts were measured in 5 accessions, 3 carrying *KCS4*(Met) and two carrying *KCS4*(Thr), and five allelic mutants from those 5 accessions. Plants were grown in control conditions (CHD) or subjected to heat and darkness for 24 h (HD) and harvested both group at the same time point. Three biological replicates were used for each line, results from separate accessions were pooled according to the allele carried. Fold change of expression was obtained using the 2^-(ΔΔCT)^ method using GAPDH3 (At1g13440) as housekeeping gene. Statistical significance was performed with ANOVA followed by Fisher LSD. Means with a common letter are not significantly different (P > 0.05).

**Supplementary Figure 6. Phylogenetic signal of the lipidome**

**a**. Distribution of the phylogenetic signal calculated for all measured lipids in HD and CHD conditions. Only lipids exhibiting significant phylogenetic signal are labelled. Lipids significant in CHD and HD are marked with blue and red dots, respectively. Compounds significant in both conditions (p<0.05) have overlapping red and blue markers. **b**. K parameter distribution obtained for the TAG profiles in CHD (control) and HD (treatment) condition according to Blomberg et al. (2003).

**Supplementary Figure 7. Prediction of climate parameters from lipidomic profiles**

Regression coefficients of selected predictive lipidomic features towards 19 climate variables and 3 geographical coordinates (latitude, longitude and altitude) of the geographical origin of analyzed accessions. Statistical evaluation of the fitted models is provided on the right-hand panel. The model fit significance (red asterisks) is estimated by the correlation coefficient between the original and model-fitted variable values. The significance of Cross-Validation test (black asterisks) represents the significance of the correlation coefficient between the predicted and real values of the test sample. The cross-validation has been performed 100 times with the 3:1 proportion of the training to test sample set size.

**Supplementary File 1**.

Alignment used for the generation of the SNP matrix used in TASSEL to analyze more polymorphisms in the QTL associated with *KCS4*. Sequences were obtained from the Arabidopsis 1001 genome project (http://signal.salk.edu/atg1001/3.0/gebrowser.php, 162 accessions). TAIR10 positions of the extract are chromosome 1: 6689119 - 6769118. Alignment was performed using MAFFT (Katoh et al., 2019) and manually curated when needed.

**Supplementary File 2**.

Alignment of KCS4 sequences from wild type and knockout accessions used in this study.

**Supplementary Dataset 1**.

Phenotypic data used for GWAS and collected in: (1) control conditions for the heat and dark stress treatment (CHD), (2) heat and dark stress (HD), (3) control conditions for the dark treatment (CD), (4) three days extended darkness (3D) and (5) six days extended darkness. Each phenotype is in different excel sheets. Plants were grown and harvested as described in Materials and Methods. Lipid classes detected are DAG: dyacylglicerols, DGDG: digalactosyldiacylglycerol, MGDG: monogalactosyldiacylglycerol, PC: phosphatidylcholine (lecithin), PE: phosphatidylethanolamine and TAG: triacylglycerol. For each species the number of carbons and the degree of bond-saturation is stated. Phenotypic value is the logarithm in the second base of intensity obtained in LC-MS.

**Supplementary Dataset 2**.

Table with GWAS results obtained from GAPIT for each phenotype presented in Supp. Table 1. All SNPs with a LOD score > 5.3 for all traits obtained in the same condition are summarized in the same table. Each sheet presents the results for each condition (see Supp. Table 1). MAF: minor allele frequency.

**Supplementary Dataset 3**.

Full list of GWAS results summarized in number of QTL (quatitative trait loci) and genes detected per condition, for all lipid species phenotyped (LOD score >5.3). Co-localization between traits in the same conditions are stated in “Peak.ID” columns. Co-localizing genes between CHD and HD condition are also mentioned. Putative lipid functions for genes related to lipid biosynthesis pathways are included in the last column. KCS4 association is highlighted in red. Each sheet compiles results for all traits phenotyped in one condition (see Supp. Table 1)

**Supplementary Dataset 4**.

Data used for TASSEL analysis: Genotypic matrix, generated from the alignment in Supp. File 1; principal components and kinship matrix used to control for population structure and confounding effects. Phenotypic matrixes for HD and 3D are included in Supplementary Dataset 1.

**Supplementary Dataset 5**.

GWAS results for HD and 3D data (see Supp. Table 1 for phenotype matrix and Supp. Table 4 for genotype matrix) using complete sequence information for 80 kb around KCS4. SNPs super-passing Bonferroni-corrected p-values (LOD > 4.38) are marked in bold. Markers ID in Tassel are referenced to positions in Supp. File 1, marker ID in GAPIT and TAIR10 position.

**Supplementary Dataset 6**.

All lipids measured in Arabidopsis wild-type accessions and *kcs4* allelic mutants. Accession ID (common name), Type (wild type or mutant), KCS4_allele (Methionine or Threonine), Condition (control or HD) are included in different columns. The table presents trait values for each biological replicate for accessions separately. Values in Figure 4 were obtained by averaging accessions encoding for a Methionine or Threonine, respectively for each type (WT and KO).

**Supplementary Dataset 7**.

Primers and oligonucleotide sequences used in this study.

