## Supplemental Figures for "Hello darkness, my old friend: 3-Ketoacyl-Coenzyme A Synthase4 is a branch point in the regulation of triacylglycerol synthesis in *Arabidopsis thaliana*"

Supp. Figure 1

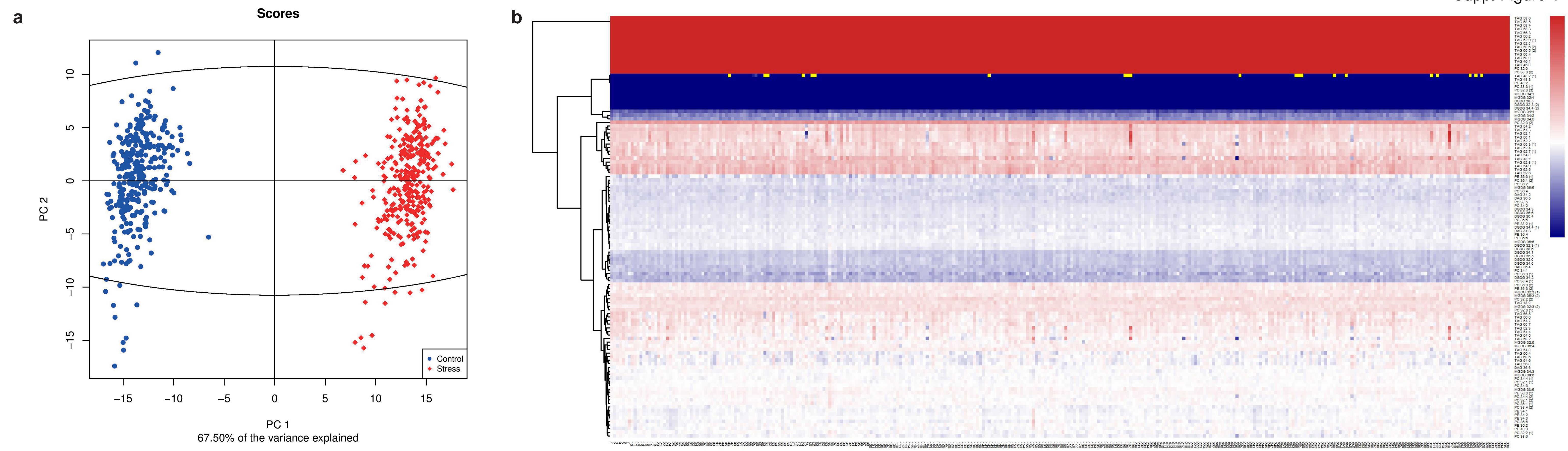


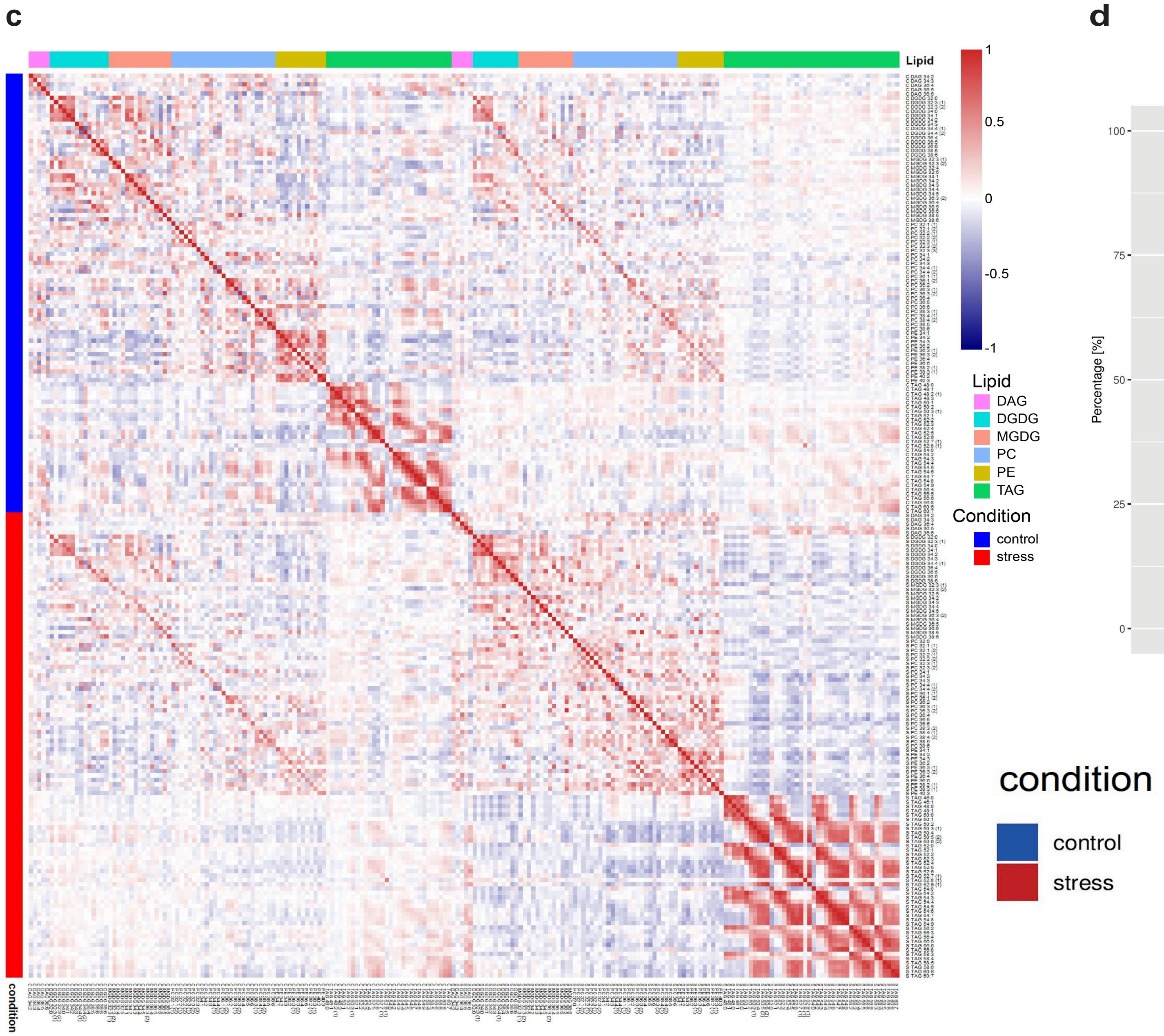


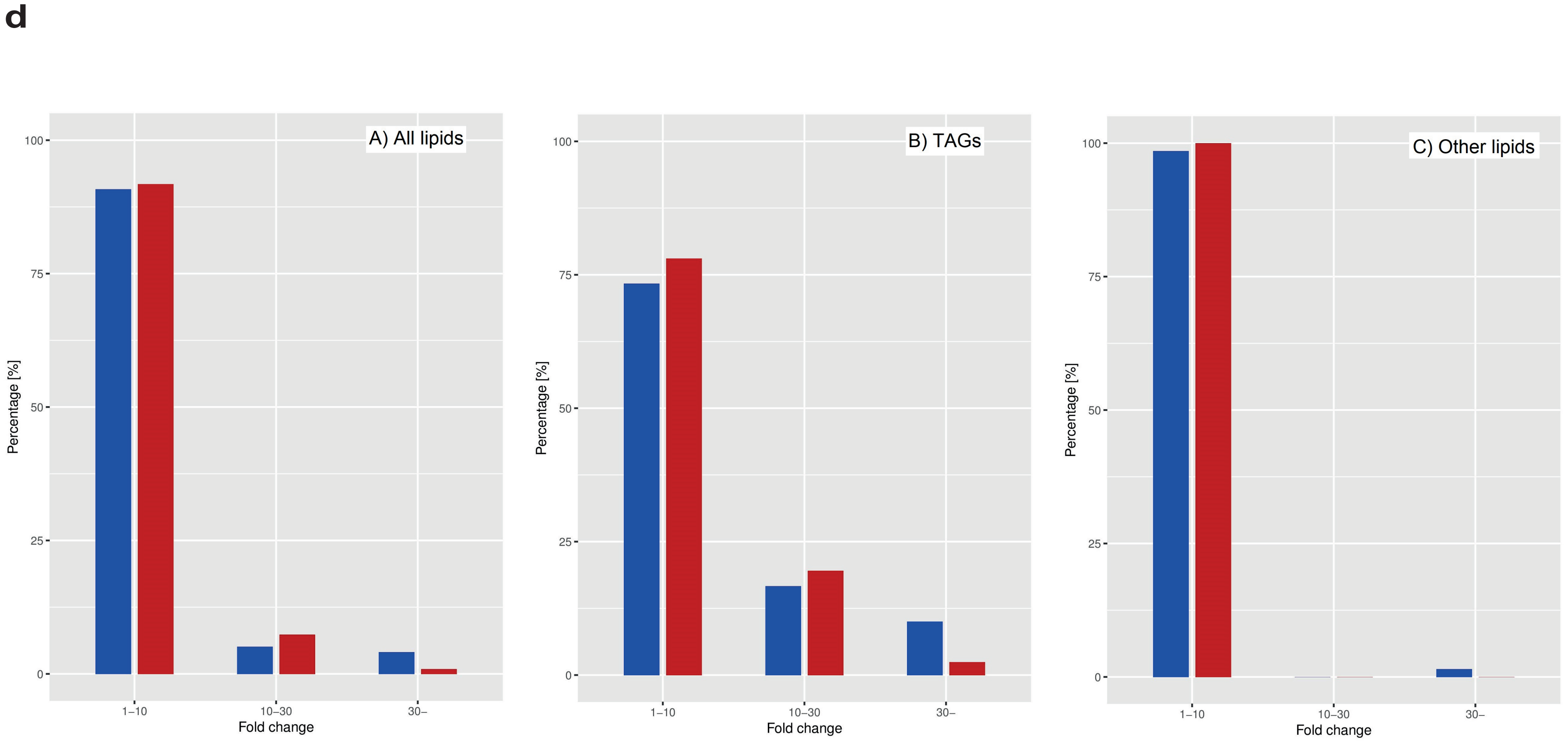


**Supplementary Figure 1. Phenotypic analyses for the Arabidopsis GWAS panel under control and stress conditions**
**a.** Principal component analysis (PCA) of lipid data for control condition (CHD, blue circles) and heat and dark condition (HD, red diamonds). Each dot represents a single accession. PC1 and PC2 explain together 67.5% of the variance observed in the phenotypic data. **b.** Heatmap of differential accumulation of lipid levels in stress (HD) and control (CHD) conditions. Lipids are clustered by lipid class. For each accession, the lipid level in HD was subtracted by the lipid level in CHD. Positive values (red) denote higher levels of lipid in HD condition, negative values (blue) denote higher level of lipid in CHD condition (scale= log2 (  of lipid intensity). **c.** Correlation pattern among lipid levels in both conditions: CHD (blue rectangle) and HD (red rectangle). Pairwise Pearson correlations (r2) were calculated between each lipid across all accessions. Lipids are highlighted with colors according to lipid classes (references included in the figure). **d.** Distribution of fold change of lipid–feature levels across all accessions. For each lipid feature, fold change was calculated by dividing the average of the five highest intensities measured in a given 5 accession by the average of the five lowest intensities. Y-axes is expressed in percentage of (A) all metabolite features (N=109 in stress, N=98 in CHD), (B) only TAG features (N=41 in stress, N= 30 in CHD) and (C) other lipid features (N=68 in stress and N=68 in CHD).

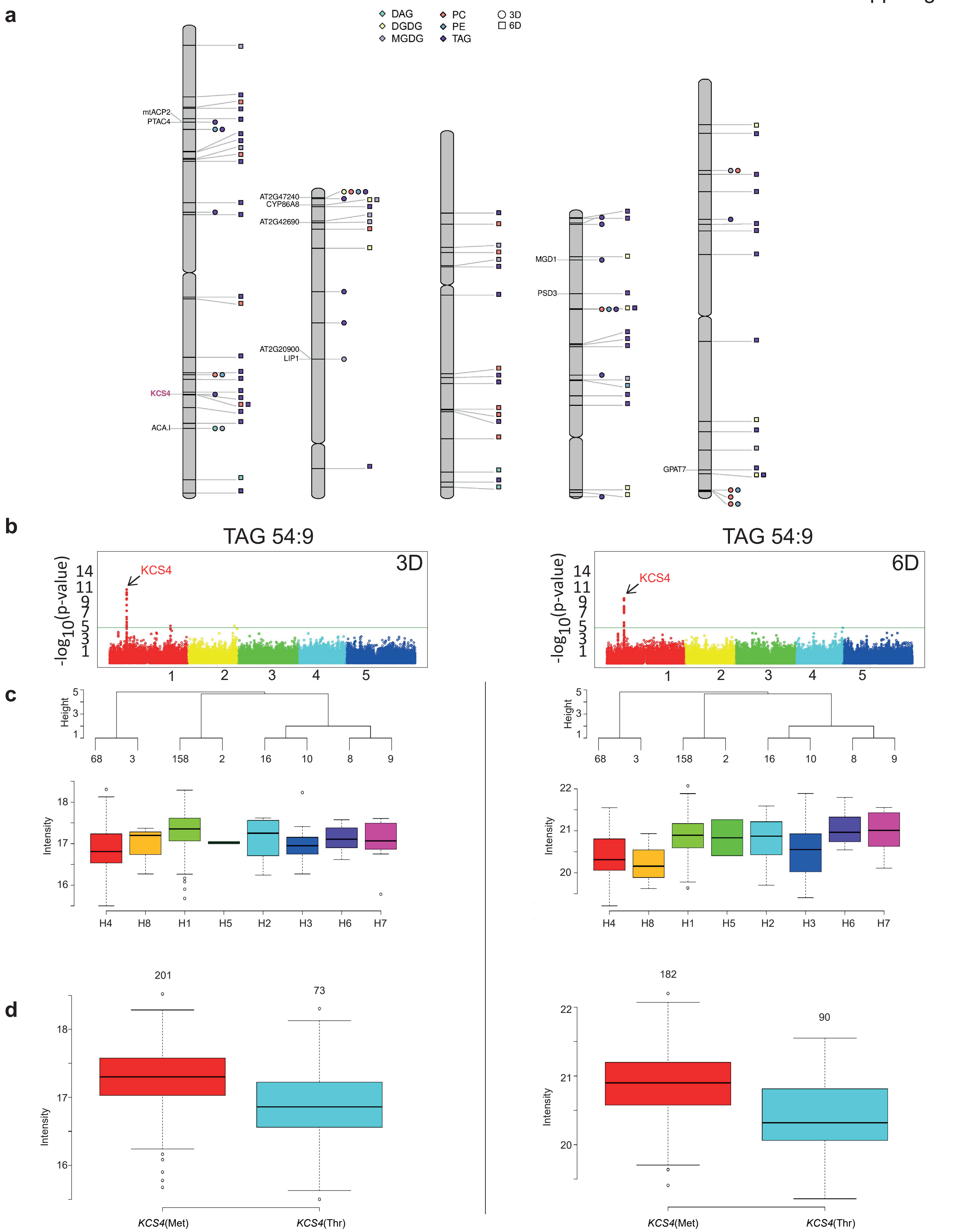


**Supplementary Figure 2. Results from GWAS analysis on lipidomic data from plants grown under extended darkness (3D, 6D) conditions.**

**a.** Chromosome scheme and Quantitative Trait Loci (QTL) identified in the two experimental conditions (3D = circles, 6D = squares), for different lipid classes: phosphatidylethanolamine (PE), phosphatidylcholine (lecithin) (PC), monogalactosyldiacylglycerol (MGDG), digalactosyldiacylglycerol (DGDG), diacylglycerol (DAG) and triacylglycerol (TAG). Color/shape references are included in the figure. Gene IDs are included only for QTL co-localizing for two or more lipid species with LOD score > 5.3. **b.** Manhattan plots obtained for TAG 54:9 in 3D (left) and in 6D (right). **c.** Average trait value (intensity of TAG 54:9, log2 scale) for the different *KCS4* haplotypes using SNPs m11502, m11503, m11504 and m11505. **d.** TAG 54:9 average value for accessions carrying the *KCS4*(Met) and *KCS4*(Thr) alleles.

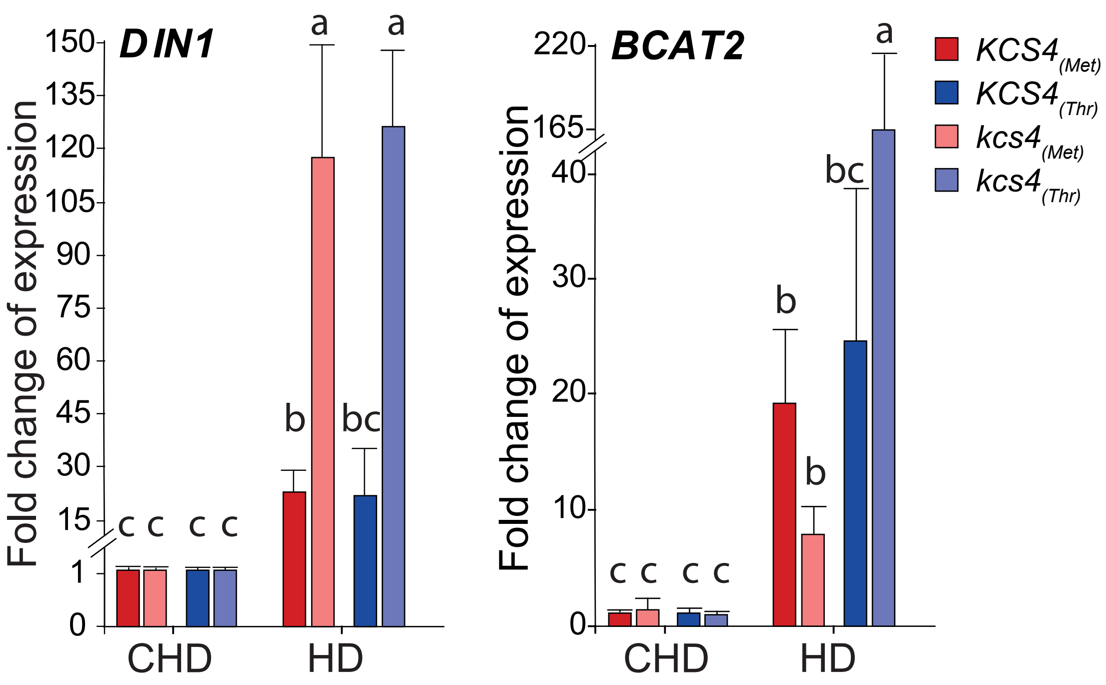


**Supplementary Figure 3. Quantification of starvation markers in *KCS4* wild-type accessions and allelic mutants.**

Transcripts were measured in 5 accessions, 3 carrying *KCS4*(Met) and two carrying *KCS4*(Thr), and five allelic mutants from those 5 accessions. Plants were grown in control conditions (CHD) or subjected to heat and darkness for 24 h (HD) and harvested both group at the same time point. Three biological replicates were use for each line, results from separate accessions were pooled according to the allele carried. Fold change of expression was obtained using the $2^{-(\Delta\Delta CT)}$ method using *GAPDH3* (At1g13440) as housekeeping gene. Statistical significance was performed with *ANOVA* followed by Fisher LSD. Means with a common letter are not significantly different (P > 0.05).

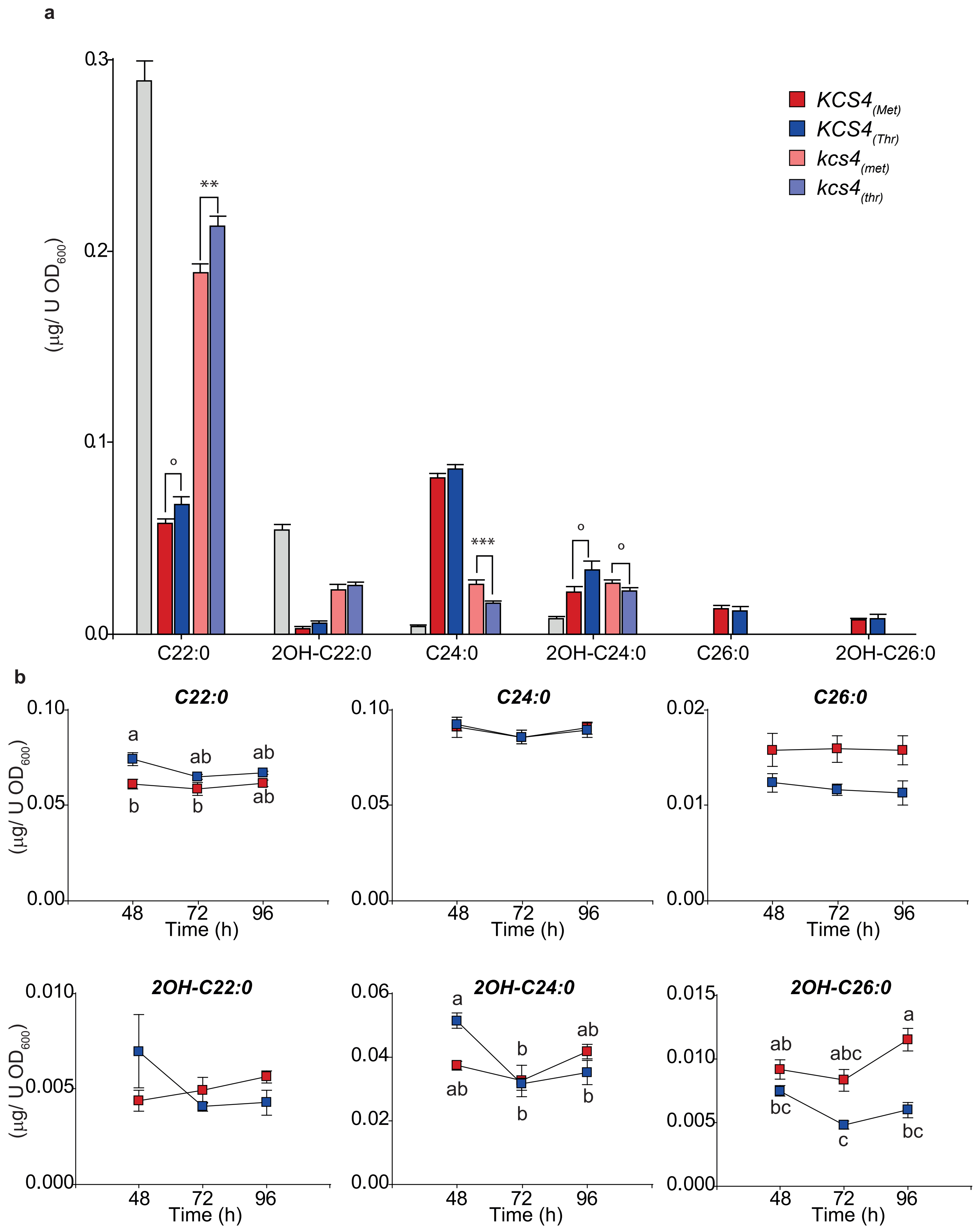


**Supplementary Figure 4. KCS4 enzymatic activity.**
**a.** Very long chain fatty acid methyl ester (VLC-FAME) profiles (22 to 26) of Δelo3 yeast mutant transformed with wild-type At*KCS4*(Met)/At*KCS4*(Thr) and allelic mutants *kcs4*(Met)/*kcs4*(Thr). References for colors are in the figure. Significant differences between alleles obtained with *t*-Test: ° $P < 0.1$, ** $P < 0.01$, *** $P < 0.001$). **b.** VLC-FAME profiles (22 to 26) at different times of harvest (48, 72 and 96 h) for At*KCS4*(Met) and At*KCS4*(Thr) wild-type alleles. Significant differences were obtained with ANOVA followed by Tukey test ($P < 0.05$).

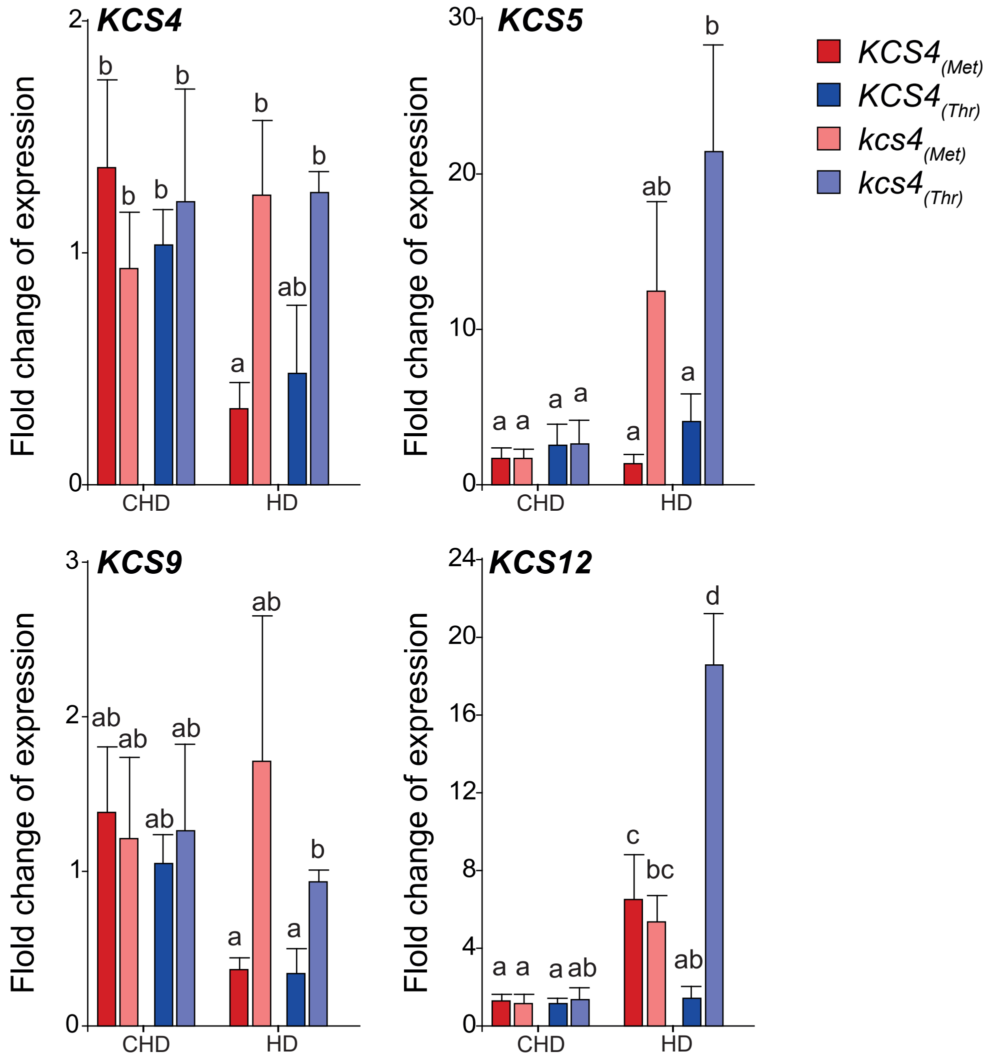


**Supplementary Figure 5. Quantification of *KCS4*, *KCS5*, *KCS9* and *KCS12* transcripts in wild-type *KCS4*(Met)/*KCS4*(Thr) accessions and *kcs4*(Met)/*kcs4*(Thr) allelic mutants.**

Transcripts were measured in 5 accessions, 3 carrying *KCS4*(Met) and two carrying *KCS4*(Thr), and five allelic mutants from those 5 accessions. Plants were grown in control conditions (CHD) or subjected to heat and darkness for 24 h (HD) and harvested both group at the same time point. Three biological replicates were used for each line, results from separate accessions were pooled according to the allele carried. Fold change of expression was obtained using the $2^{-(\Delta\Delta CT)}$ method using *GAPDH3* (At1g13440) as housekeeping gene. Statistical significance was performed with *ANOVA* followed by Fisher LSD. Means with a common letter are not significantly different ($P > 0.05$).

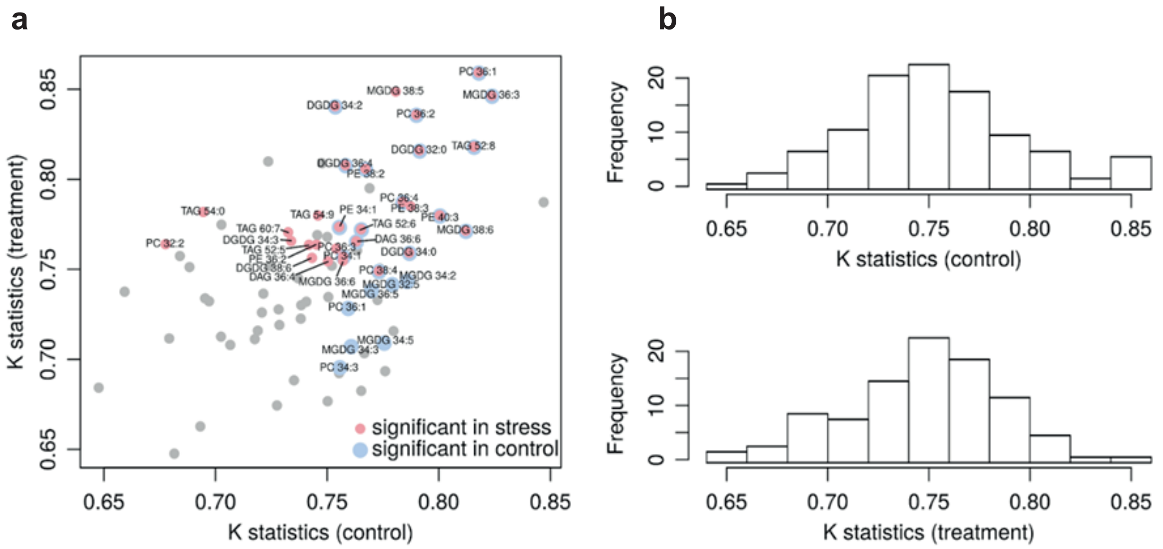


**Supplementary Figure 6. Phylogenetic signal of the lipidome**

**a.** Distribution of the phylogenetic signal calculated for all measured lipids in HD and CHD conditions. Only lipids exhibiting significant phylogenetic signal are labelled. Lipids significant in CHD and HD are marked with blue and red dots, respectively. Compounds significant in both conditions ($P < 0.05$) have overlapping red and blue markers. **b.** K parameter distribution obtained for the TAG profiles in CHD (control) and HD (treatment) condition according to (Blomberg et al., 2003).

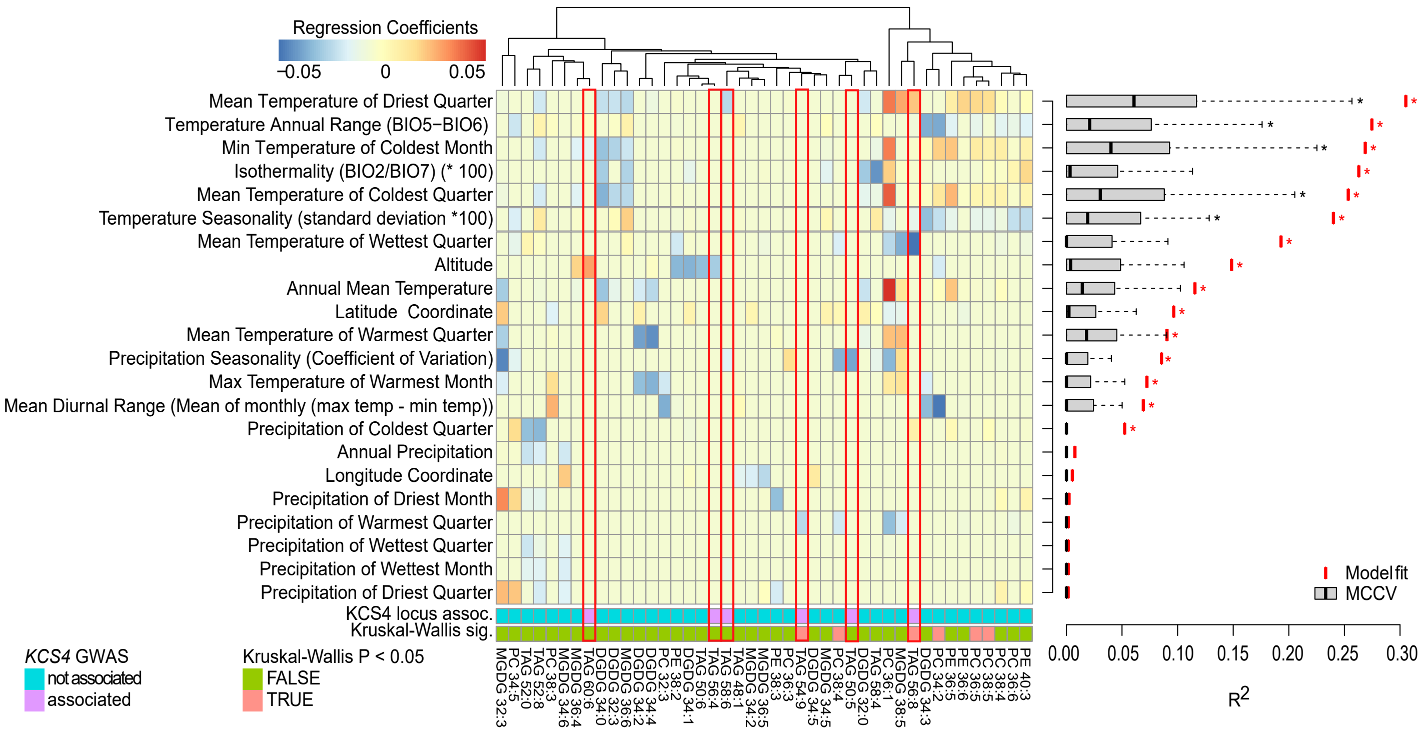


**Supplementary Figure 7. Prediction of climate parameters from lipidomic profiles**

Regression coefficients of selected predictive lipidomic features towards 19 climate variables and 3 geographical coordinates (latitude, longitude and altitude) of the geographical origin of analyzed accessions. Statistical evaluation of the fitted models is provided on the right-hand panel. The model fit significance (red asterisks) is estimated by the correlation coefficient between the original and model-fitted variable values. The significance of Cross-Validation test (black asterisks) represents the significance of the correlation coefficient between the predicted and real values of the test sample. The cross-validation has been performed 100 times with the 3:1 proportion of the training to test sample set size.
